# De novo detection of somatic variants in high-quality long-read single-cell RNA sequencing data

**DOI:** 10.1101/2024.03.06.583775

**Authors:** Arthur Dondi, Nico Borgsmüller, Pedro F. Ferreira, Brian J. Haas, Francis Jacob, Viola Heinzelmann-Schwarz, Tumor Profiler Consortium, Niko Beerenwinkel

**Affiliations:** ETH Zurich, Department of Biosystems Science and Engineering, Schanzenstrasse 44, 4056 Basel, Switzerland; SIB Swiss Institute of Bioinformatics, Schanzenstrasse 44, 4056 Basel, Switzerland; Broad Institute of Massachusetts Institute of Technology and Harvard, Cambridge, Massachusetts, USA; University Hospital Basel and University of Basel, Department of Biomedicine, Ovarian Cancer Research, Hebelstrasse 20, 4031 Basel, Switzerland

## Abstract

In cancer, genetic and transcriptomic variations generate clonal heterogeneity, leading to treatment resistance. Long-read single-cell RNA sequencing (LR scRNA-seq) has the potential to detect genetic and transcriptomic variations simultaneously. Here, we present LongSom, a computational workflow leveraging high-quality LR scRNA-seq data to call *de novo* somatic single-nucleotide variants (SNVs), including in mitochondria (mtSNVs), copy-number alterations (CNAs), and gene fusions, to reconstruct the tumor clonal heterogeneity. Before somatic variants calling, LongSom re-annotates marker gene based cell types using cell mutational profiles. LongSom distinguishes somatic SNVs from noise and germline polymorphisms by applying an extensive set of hard filters and statistical tests. Applying LongSom to human ovarian cancer samples, we detected clinically relevant somatic SNVs that were validated against matched DNA samples. Leveraging somatic SNVs and fusions, LongSom found subclones with different predicted treatment outcomes. In summary, LongSom enables *de novo* variants detection without the need for normal samples, facilitating the study of cancer evolution, clonal heterogeneity, and treatment resistance.

## Introduction

Cancer cells accumulate somatic genomic variations, such as single-nucleotide variants (SNVs), copy number alterations (CNAs), and gene fusions during their lifetime, leading to intratumor heterogeneity, i.e., the existence of cancer subpopulations with distinct genotypes and phenotypes. This is presumed to be a leading cause of therapy resistance and one of the main reasons for poor overall survival in cancer patients with metastatic disease (Jamal-Hanjani et al. 2015; Vasan et al. 2019; Ramón Y Cajal et al. 2020). While genetic mechanisms are often evoked as the primary cause of therapeutic resistance in those subpopulations, the adaptive mechanisms underlying therapy resistance are both of genetic (SNVs, CNAs, gene fusions, etc.) and non-genetic (epigenetic, transcriptomic, microenvironment, etc.) origin (Mansoori et al. 2017; Marine et al. 2020). The first step in identifying therapy-resistant subclones is, therefore, to examine these interlinked features jointly (Foord et al. 2023) by capturing genetic and transcriptomic variants at the single-cell level (Mansoori et al. 2017; Dagogo-Jack and Shaw 2018; Marine et al. 2020).

Droplet-based scRNA-seq (e.g. 10X Genomics Chromium) can detect same-cell genetic and transcriptomic variants. However, those protocols can only capture RNA molecules via their 3’ or 5’ ends, and short-read (SR) scRNA-seq coverage is heavily biased towards the 3’/5 end of genes. Recently, methods to call SNVs (Zhang et al. 2023; Muyas et al. 2024) and CNAs (Serin Harmanci et al. 2020), (Gao et al. 2021, 2023) in SR scRNA-seq were developed, compensating the 3’ capture bias by pooling large amounts of cells or sequencing at very high read depths. However, SR scRNA-seq is unsuited to detect isoforms or gene fusions. Long-read (LR) scRNA-seq, in contrast, sequences full-length RNA molecules, and we have shown in recent work that high-quality LR scRNA-seq can simultaneously detect clinically relevant SNVs, CNAs, fusions, and isoform-level expression in the same cells (Joglekar et al. 2021; Dondi et al. 2023; Shiau et al. 2023; Al’Khafaji et al. 2024; Qin et al. 2024).

Somatic variants are typically identified by comparing variants from tumor biopsies with those from matched normal biopsies derived from respective healthy tissue. However, as matched normals are rarely available, methods to detect somatic variants de novo were developed for bulk sequencing, mainly relying on germline VAF profiles and tumor purity estimates combined with extensive filtering against public databases (Teer et al. 2017; Sun et al. 2018; Shiau et al. 2023). As sensitivity/recall is similar to variant detection with a matched normal, but precision is lower, tumor-only detection in bulk sequencing is usually deemed more appropriate for detecting known mutations than detecting variants de novo (Teer et al. 2017). For scRNA-seq, SComatic (Muyas et al. 2024) was developed to call variants de novo without matched DNA-seq normal, leveraging non-cancer microenvironment cells in the tumor biopsy to differentiate somatic from germline variants. This approach relies on initial cell type annotation based on gene expression patterns, an open challenge due to overlapping, poorly expressed, or incomplete marker gene sets, and even a low percentage of cancer cells misannotated as non-cancer will lead to false-negative variants filtered out as germline (Muyas et al. 2024). Consequently, as somatic variant calling depends intrinsically on the quality of cell type annotations, methods ensuring the correctness of the annotations are needed.

Cell types and clonal substructures are traditionally identified in scRNA-seq using gene expression profiles. However, identifying different cancer clones requires sufficient transcriptional divergence between them. Instead, clonal substructures can also be reconstructed using somatic variants (Zhou et al. 2020; Gao et al. 2021, 2023; Kannan et al. 2022; Muyas et al. 2024). For this, mitochondrial SNVs (mtSNVs) serve as an excellent complement to nuclear SNVs and fusions (Kwok et al. 2022; Miller et al. 2022), as mitochondrial RNA (mtRNA) is highly available (Osorio and Cai 2021) and mutated (>10-fold higher than in the nuclear genome (Wallace 1994)) in scRNA-seq. mtSNVs can also be pathogenic in cancer (Koshikawa et al. 2017), yet, few variant calling methods properly integrate mtSNVs detection (Mukherjee et al. 2023).

This study aims to provide a computational workflow for detecting variants (SNVs, mtSNVs, fusions, and CNAs) in LR scRNA-seq of tumor tissue samples without requiring matched-normal sample, subsequently integrating them to reconstruct the samples’ clonal heterogeneity, and identifying subclones with different predicted treatment outcomes.

## Results

### Overview of LongSom, a computational workflow for LR scRNA-seq variants detection and clonal reconstruction

We developed LongSom, a workflow for detecting genetic variants and finding cancer subclones in LR scRNA-seq data without requiring matched normal. Briefly, LongSom takes BAM files and cell type annotations as input, reassesses the cell type annotations, subsequently calls SNVs, mtSNVs, fusions, and CNAs in single-cells based on the re-annotated cell types, and finally reconstructs the clonal heterogeneity (**Figure 1a**).

**Figure 1:**
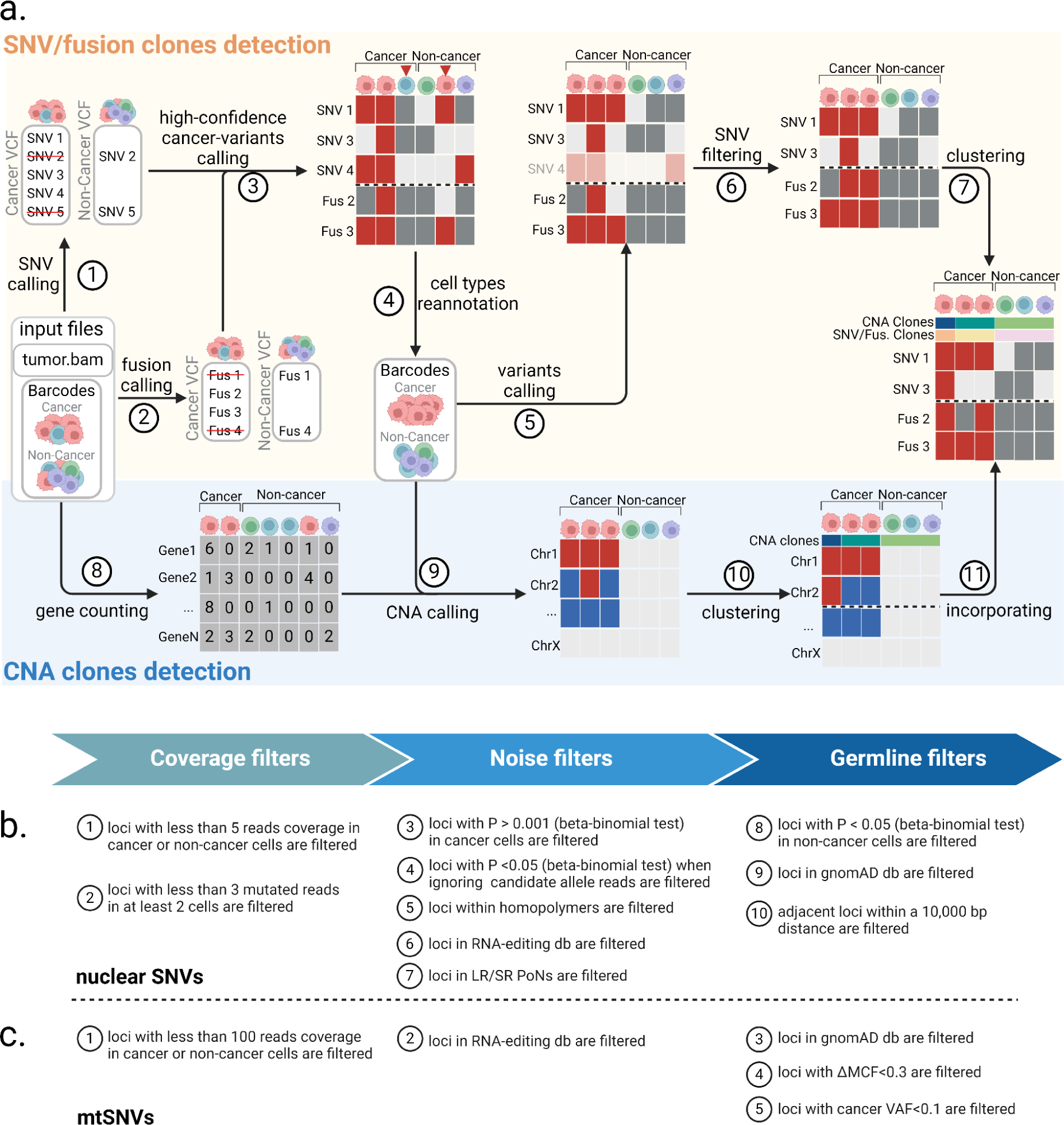
Overview of LongSom. **a.** LongSom’s methodology for detecting somatic SNVs, fusions, and CNAs and subsequently inferring cancer subclones in LR scRNA-seq individual patients data. (1) SNV and (2) fusion candidates are detected from pseudo-bulk samples. (3) High-confidence cancer variants (SNVs and fusions) are selected based on mutated cell fraction in cancer and non-cancer cells. (4) Cells are re-annotated based on high-confidence cancer variants. (5) A new set of candidate variants is called based on re-annotated barcodes. (6) Candidate SNVs are filtered through a set of 10 filters. (7) cells are clustered based on somatic fusions and SNVs. In parallel, (8) gene expression per cell is computed, (9) CNAs are detected, (10) cells are clustered based on CNAs, and (11) CNA clones are incorporated to the fusions and SNVs clustered matrix. **b.** Candidate nuclear SNV successive filtering steps. Candidates passing all 10 steps are called as somatic SNVs (Methods). **c.** Candidate mtSNVs filtering steps. ΔMCF represents the difference of mutated cells fraction between cancer and non-cancer cells. Candidates passing all 5 steps are called as somatic mtSNVs (Methods).

To avoid false-negative calls due to cell type misannotation or cells containing high levels of ambient cancer RNA, LongSom re-annotates the marker-based cell types. For this, LongSom calls a set of ‘high-confidence cancer variants’ (SNVs, mtSNVs, and fusions) following eight filtering steps, and reannotates cells based on their mutational burden (Methods). LongSom then re-calls variants using reannotated cell types. Somatic SNVs are identified in ten filtering steps including hard filters and statistical testing (**Figure 1b**, Methods). Calling mtSNVs remains challenging, as high levels of ambient mtRNA, released by dead or dying cells (Young and Behjati 2020), can cause false negative calls on the bulk level (loci wrongly excluded as germline) and false positive calls on the single-cell level (contaminated non-cancer cells called as mutated). Therefore, LongSom treats mtSNVs differently from nuclear SNVs and calls them in five filtering steps (**Figure 1c**, Methods). Finally, LongSom infers the clonal structure of the samples using two different approaches. One approach leverages the detected SNVs, mtSNVs and fusions as input for the Bayesian non-parametric clustering method BnpC (Borgsmüller et al. 2020). The other approach predicts CNAs based on gene expression in cancer cells and defines subclusters using inferCNV (https://github.com/broadinstitute/infercnv) (**Methods**).

### LongSom reannotates cell types based on mutational profiles

We applied LongSom to previously published high-quality (PacBio) LR scRNA-seq data of omentum metastasis samples obtained from three chemo-naive HGSOC patients: P1, P2, and P3 (Dondi et al. 2023). Those samples were composed of 337 cancer cells (41,959 median UMI per cell) and 1225 micro-environment cells (11,716 median UMI per cell), referred as “non-cancer cells” in the following. After cell-type reannotation, we found that cells reannotated as cancer were mostly clustering with cells previously annotated as cancer cells based on expression data, but some clustered with non-cancer cells (**Figure 2a**). We found that 8, 2, and 27% of the cells that LongSom annotated as cancer were previously annotated as non-cancer cells in the tumor biopsy samples of patients P1, P2, and P3, respectively (**Figure 2b**). The tumor biopsy of patient P3 had only 10% cancer cells (Dondi et al. 2023), which could explain the high level of cell misannotation. Cells reannotated from cancer to non-cancer cells had a similar mutational burden than cells previously annotated as non-cancer in all patients. In patients P1 and P2, cells reannotated from non-cancer to cancer cells had a similar mutational burden and mean fraction of mutated loci as cells previously annotated as cancer (P>0.05, Tukey-Krammer’s test (Kramer 1956)), while being significantly different from cells previously annotated a non-cancer (P<0.001) **(Figure 2c,d**). In P3, cells reannotated from non-cancer to cancer cells were significantly different from cells previously annotated as both cancer and non-cancer. Those cells were likely misannotated due to low expression in the first place, leading to a lower mutational burden. Furthermore, cells with the lowest mutational burden are also cells not clustering with cancer cells (**Figure 2a, Supplementary Figure S1a**). Those are likely cells with high levels of ambient RNA, and while likely not cancer cells, they would still cause false negatives if they were not reannotated. In summary, cell type reannotation reduced the cell-variants noise (**Figure 2e, Supplementary Figure S1b-c**), and in the following, cancer or non-cancer cells refer to the reannotated cell types.

**Figure 2:**
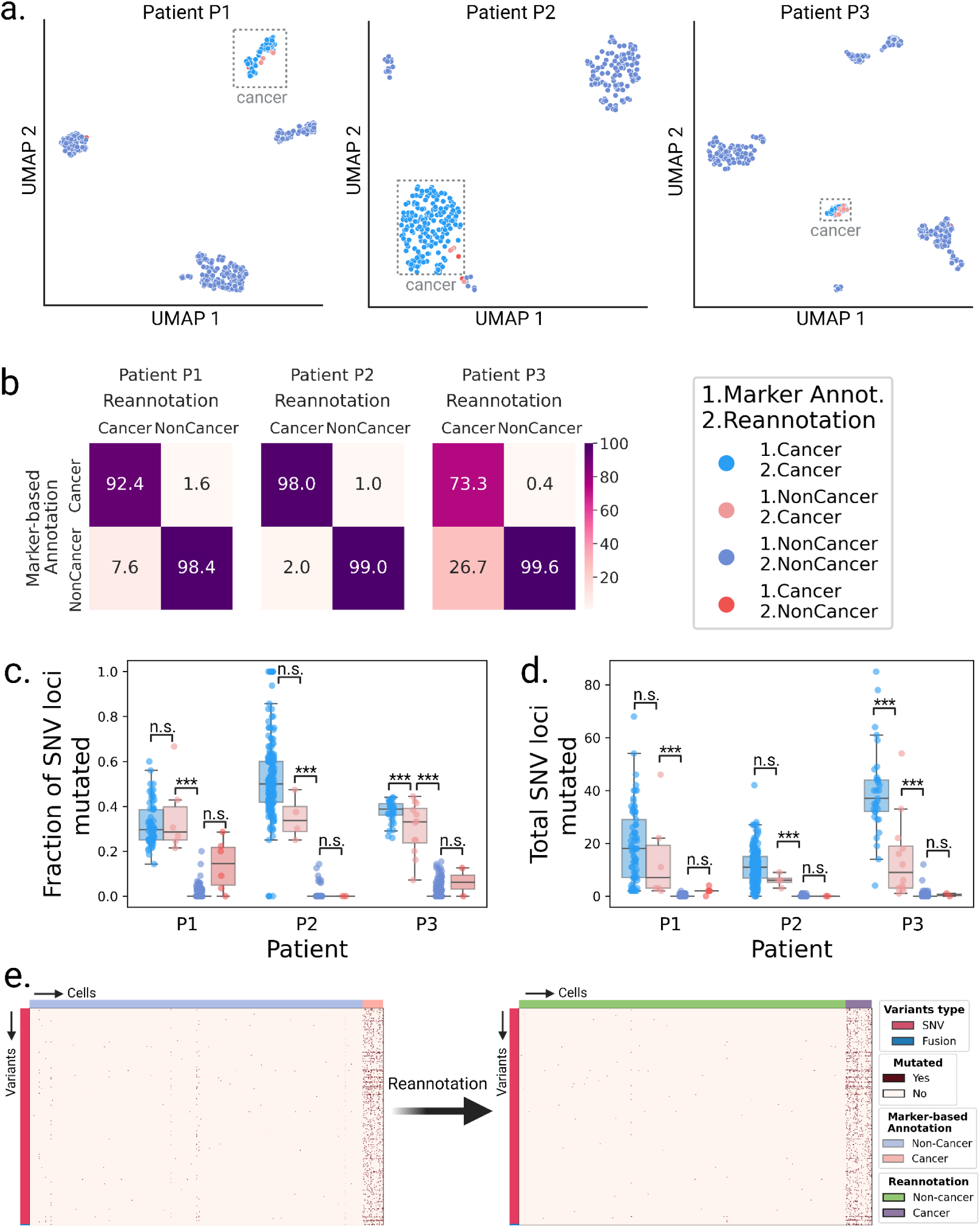
Validation of the cell type re-annotation based on mutational profiles. **a.** UMAP embeddings of LR scRNA-seq expression per patient. Cells are colored by annotation status; light-red cells were previously predicted as non-cancer using marker gene expression-based annotation and were reannotated as cancer by LongSom based on high-confidence cancer variants.**b.** Confusion matrices of cells predicted as cancer or non-cancer using marker genes, and cells reannotated as cancer or non-cancer by LongSom, colored and annotated by the percentage of the total number of cells in each category. E.g. the bottom left square represents cells previously annotated as non-cancer that were reannotated as cancer (false negative cancer cells). **c,d** Boxplots of **c.** the fraction of SNV loci that were found mutated in each cell, considering only loci with minimum coverage of 1 read at the locus in a cell, and **d.** the total number of SNVs mutated in each cell, per patient, colored by their annotation status. Points represent individual cells, and boxes display the first to third quartile with median as horizontal line, whiskers encompass 1.5 times the interquartile range. P values were calculated using Tukey-Krammer’s test and are described with the following symbols: n.s : P > 0.05, *: P ≤ 0.05, **: P ≤ 0.01, ***: P ≤ 0.001. **e.** Cell-variant matrixes of patient P3 before (left) and after (right) re-annotation. Red indicates that a locus is mutated in a cell (bet-binomial test P value < 0.05), and white that it is not (either P > 0.05 or no coverage).

### Validation of LongSom somatic calls using scRNA-seq and scWGS data

After reannotation, we processed 4,271,449 candidate loci with at least one mutated read and five reads coverage in aggregated cancer cells (**Figure 3a**). LongSom called 822 somatic SNVs passing all filters, which mapped to intronic regions (75%), exonic regions (10.9%), 3’UTR regions (6.2%), intergenic regions (4.8%), splicing sites (1.7%), and 5’UTR regions (1.4%) (**Figure 3a,b**). In this dataset, we previously identified that 32% of the reads resulted from contaminating DNA internally primed on their intronic polyA-rich regions (Dondi et al. 2023), a common phenomenon in scRNA-seq known as “intra-priming” (Verwilt et al. 2023). While those reads are removed from transcript counts, they are valuable to call SNVs in scRNA-seq (as they come from same-cell DNA) and they can explain the large fraction of intronic variants we observed. A WGS study of a cohort of 962 individuals (Morrison et al. 2013) found 58% of intergenic variants, 36% of intronic variants, and 6% of “functional” (exonic, splicing, 3’UTR, and 5’UTR and splicing) variants. This study observed a 1 to 17 ratio of functional to intronic and intergenic variants, while we observed a 1 to 5 ratio, indicating that droplet-based scRNA-seq still selects functional variants despite intra-priming.

**Figure 3:**
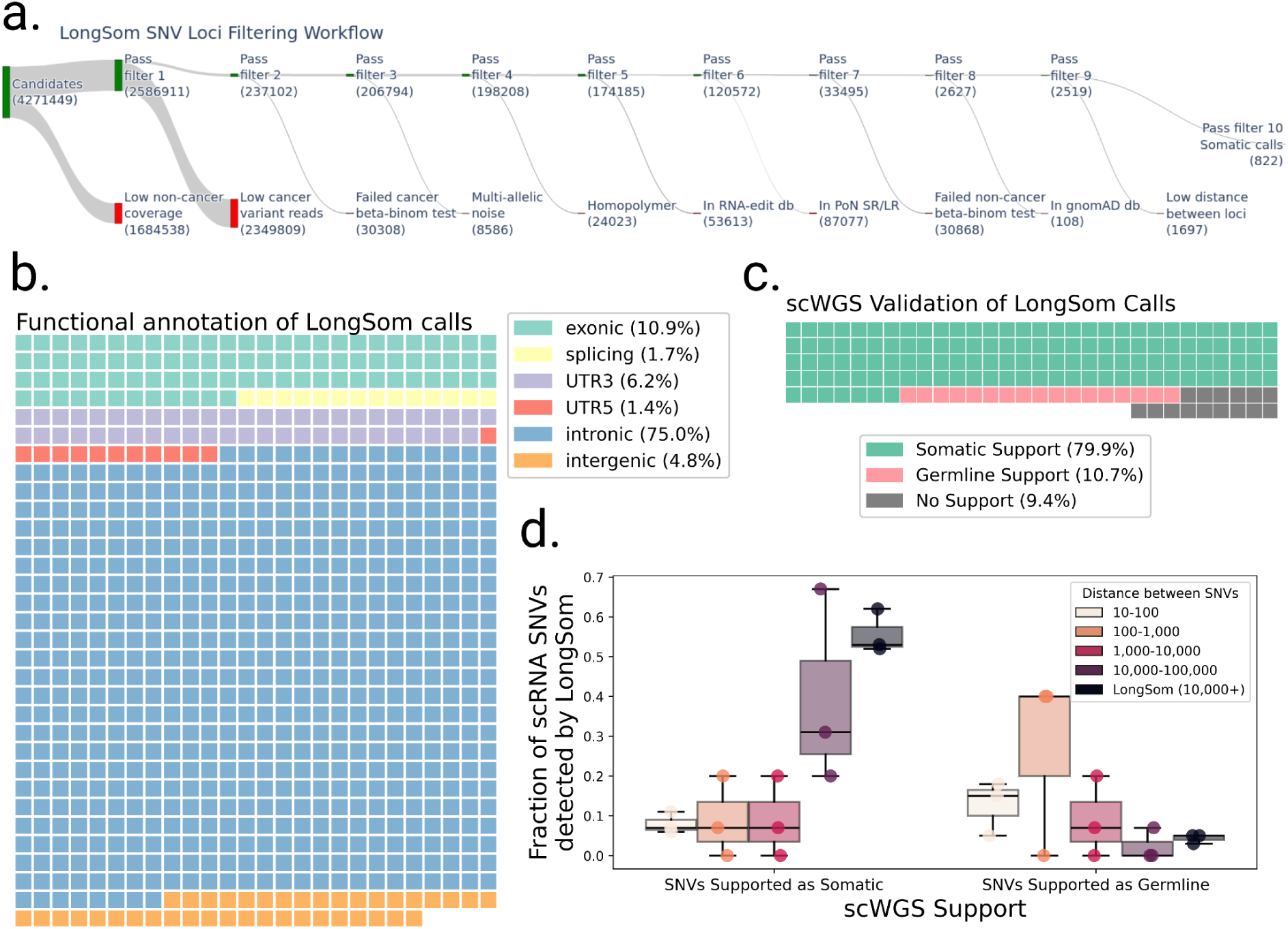
Validation of LongSom somatic calls using scRNA and scWGS data. **a.** LongSom SNV filtering workflow, indicating the number of loci passing each of the 10 filtering steps (top, green) or being filtered (bottom, red), starting from all loci with at least one mutated read and 5 reads coverage in cancer cells. **b.** Waffle plot representing each of the 822 somatic SNVs detected by LongSom, colored by their RefSeq functional annotation. **c.** Waffle plot of all loci called by LongSom with sufficient power in scWGS for validation (Methods), colored by their status in scWGS. **d.** Boxplot of the fraction of all loci called after filtering step 9 that is supported by scWGS data as either somatic or germline, colored by the distance from the closest mapping SNV also detected. LongSom calls represent all 822 calls after filtering step 10 (SNVs not within 10,000bp or less from each other). Each point is a patient.

To validate those calls, we used single-cell whole-genome sequencing (scWGS) data from matched omental metastases for each patient. In addition to diploid clones (likely non-cancer), we found two aneuploid clones (likely cancer) in patient P1 scWGS data, one in P2, and two in P3 (Methods, **Supplementary Figure S2**). In the 211 loci called as somatic by LongSom with sufficient scWGS depth, we called SNVs in scWGS clones, and loci called in at least one aneuploid clone and in none of the diploid clones were defined as somatic, while loci called in diploid clones were defined as germline. We found that 80% of the calls were supported as somatic in scWGS, while 11% were supported as germline calls and therefore likely to be false positive (**Figure 3c**). We also found that 9% of the loci were not called in any clone, possibly due to tumor heterogeneity between the scWGS and scRNA-seq samples. We also investigated the scWGS support of exonic, 3’UTR, 5’UTR and splice-site variants versus intronic and intergenic variants, and found the similar results, confirming the the intronic and intergenic variants identified are not false positives (**Supplementary Figure S3**). For patients P1 and P3, we had access to matched normal scRNA-seq data from matching distal tumor-free omental tissues (Dondi et al. 2023). As a supplementary validation, we called SNVs in those samples, and found that 13% (P1) and 4% (P3) of LongSom somatic calls were mutated in the matched normal, i.e. germline false positives (**Supplementary Figure S4**). Altogether, the high support for somatic mutations in scWGS data, as well as the low amount of false positive germline calls found in both scWGS and matched normal scRNA data show LongSom ability to call somatic variants without matched normal.

We investigated the correlation between the distance separating somatic SNVs and their support in scWGS. When taking all 2,519 loci passing filter 9 (**Figure 3a**), we found that the lower the mapping distance between two somatic SNVs, the lower the support in scWGS data, with high rates of germline and low rates of somatic variants (**Figure 3d**). Even SNVs within a 1,000-10,000bp distance had low scWGS support, suggesting gene-wise allelic expression differences between cancer and non-cancer cells (**Figure 3d**). Therefore, LongSom filters loci within a 10,000bp mapping distance from each other.

### Somatic mitochondrial reads contaminate tumor microenvironment cells in scRNA-seq and scWGS data

LongSom detected five mtSNVs in patient P1 at positions 2815, 3092, 5179, 13635 and 16192, two in P2 at positions 2573 and 16065, and none in P3. In patient P1, cancer cells had a mean VAF per cell ranging from 10 to 98% for all identified loci, while non-cancer cells had a mean VAF per cell ranging from 0.1 to 3% (**Figure 4a**). Cells from a matched normal biopsy had mean VAF per cell < 0.001% at all loci, discarding germline heteroplasmies and suggesting a contamination of non-cancer cells by cancer-derived mtRNA. In Patient P2, we also observed a similar VAF profile in non-cancer cells at locus 2573 (**Figure 4b**). To assess the levels of ambient mtRNA derived from cancer cells, we computed the VAF of each mtSNV loci in all empty droplets containing no cell and only ambient RNA (Methods). We found that empty droplets had cancer-like VAF profiles, with mean VAFs 3.4 (+/-1) times higher than the mean VAF of the biopsy (**Figure 4c**). We also found a strong correlation (R=0.93) between aggregated mutated reads in empty droplets and non-cancer cells (**Figure 4d**). In contrast, the correlation between the total coverage and the number of mutated reads was weaker (R=0.84). This supports that mutated reads observed in non-cancer cells come from cancer ambient mtRNA. We investigated the mtSNVs in a matched scWGS sample and found similar contamination profiles in cells from non-cancer clones at all loci (**Figure 3e,f**). This suggests the presence of ambient cancer mtDNA, too, and possibly of entire mitochondria. In comparison, the seven mtSNVs detected by LongSom were all filtered by SComatic (Muyas et al. 2024) due to the mitochondrial noise levels described above.

**Figure 4:**
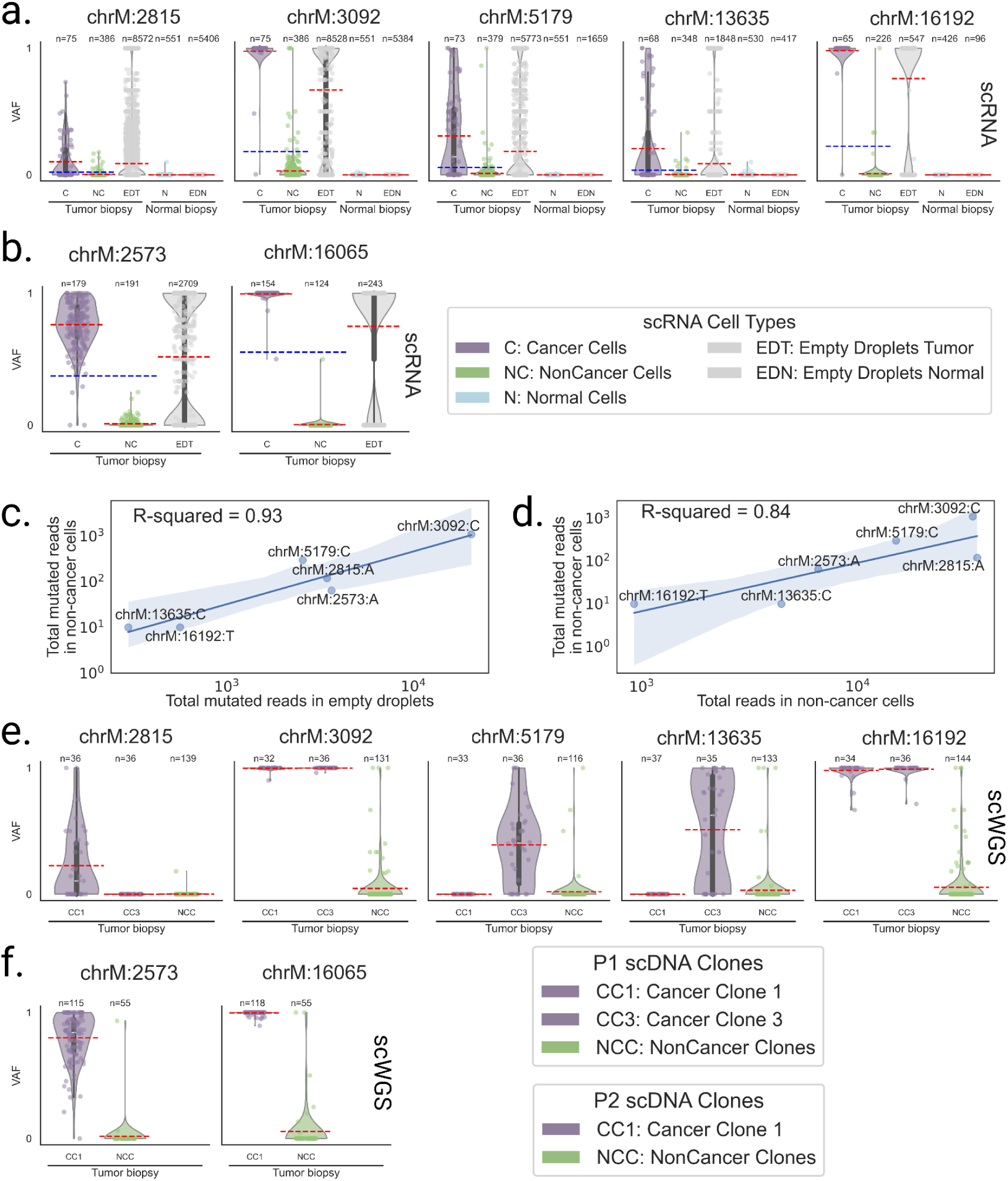
Mitochondrial reads harboring somatic mutations are detected in non-cancer cells. **a,b,** Violin plots of the VAF of each cell for **a.** patient P1 and **b.** P2 mtSNV loci in scRNA-seq data, categorized by reannotated cell types and empty droplets in tumor and normal biopsies. Individual points are cells or droplets. The blue dashed line represents the mean VAF in cells from the tumor biopsy. The red dashed line represents the mean VAF in each category. n refers to the number of cells with at least one read covering the locus. **c,d,** Log aggregated mutated reads in non-cancer cells, as a function of **c.** log aggregated mutated reads in empty droplets and **d.** log aggregated total reads in non-cancer cells, for all loci from P1 and P2 except locus 16065 in P2 which was discarded due to low expression. **e,f,** Violin plots of the VAF of each cell for **e.** patient P1 and **f.** P2 mtSNV loci in scWGS data, categorized by clones in the tumor biopsy. Individual points are cells. The red dashed line represents the mean VAF in each clone. n refers to the number of cells with at least one read covering the locus.

### LongSom outperforms SComatic for somatic variants calling

We compared somatic SNV calls from LongSom and SComatic. The main differences between the two algorithms is that LongSom corrects cell type annotations, uses a stricter beta-binomial threshold when filtering out germline variants, and filters the final calls within 10,000bp of each other. To ensure that LongSom performance was not only driven by the 10,000bp filter, we also applied this filter to SComatic calls. LongSom found 342, 145, and 340 somatic SNVs in patients P1, P2, and P3, respectively, while SComatic found 319, 155 and 260 (**Figure 5a**, **Supplementary Tables S1, S2**). The calls overlapped by only 63, 40, and 72% between the two methods in each patient, while 21, 10, and 33% of the calls were unique to LongSom, versus 16, 18, 13% for SComatic. The largest difference was observed in P3, which is concordantly the patient with the largest proportion of cells reannotated, followed by P1 (**Figure 2, 5c)**. Calls unique to LongSom had 0.82-0.83 somatic support in scWGS, similar to calls common to both methods (0.72-0.86), while calls unique to SComatic had a lower somatic support (0.38-0.50) (**Figure 5c**). Calls unique to LongSom (0-0.17) and common (0.11-0.14) to both methods also had a lower proportion of germline support in scWGS data (0.11-0.5) than calls unique to SComatic or common to both methods (**Figure 5c**).

**Figure 5:**
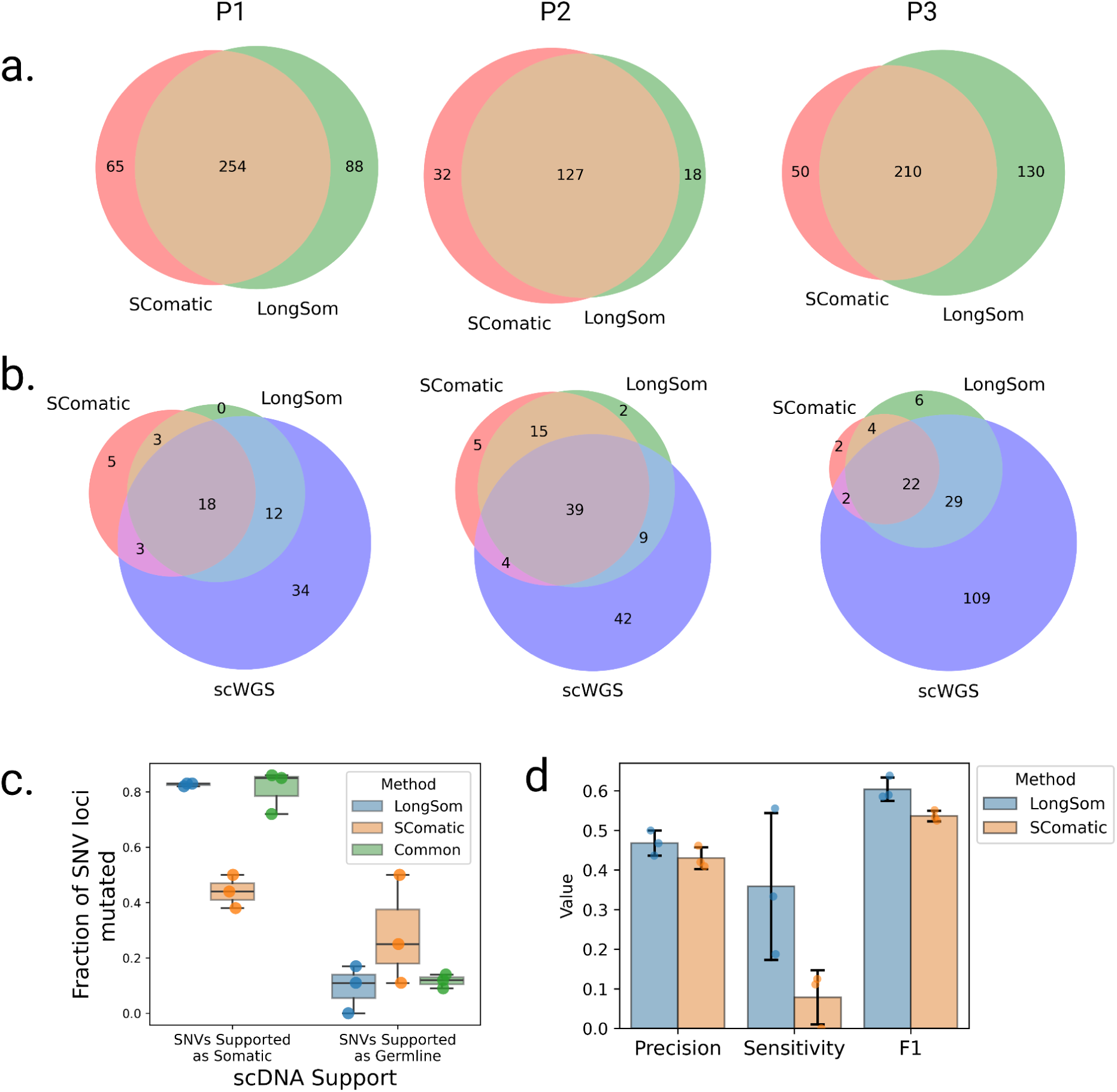
Comparison of Longsom versus SComatic performance using scWGS data. **a.** Venn diagrams of the intersection between Longsom and SComatic somatic calls in LR scRNA-seq data from patients P1, P2 and P3. **b.** Venn diagrams of the intersections between scWGS SNV calls. LongSom LR scRNA-seq calls and SComatic LR scRNA-seq calls. **c.** Boxplot of all loci called after filtering step 9, colored by the distance from the closest mapping SNV also detected, and classified by somatic or germline scWGS support. Each point is a patient. **d.** Performance of LongSom and SComatic for detecting somatic mutations in LR scRNA-seq data. Each point is a patient, the bars represent the mean value, and the error bars are the standard deviation for each statistic computed.

Next, we compared the performances of LongSom versus SComatic. To this end we considered a set of 323 mutations detected in scWGS data with sufficient coverage in scRNA-seq data for benchmarking (Methods). LongSom call set had more overlap with scWGS callset than SComatic in all patients, especially in patient P3 (**Figure 5b**). In scWGS calls supported by at least one read in scRNA-seq data, LongSom achieved a score of 0.44-0.5 precision across the three samples, similar to SComatic (0.41-0.46) (**Figure 5d**). However, LongSom achieved a superior sensitivity (0.19-0.55) than SComatic (0-0.13), and a higher F1 score. The higher sensitivity is largely due to LongSom cell reannotation that prevents false negative germline calls, whereas SComatic filters out SNVs that are supported as somatic by scWGS data.

### LongSom detects panel-validated variants

The three patients also underwent bulk panel DNA sequencing (Methods), where 29 SNVs were found (**Supplementary Table S3**). All three patients had at least one somatic SNV called in *TP53* (including a variant introducing a stop codon in patient P3) with a VAF >30%, and LongSom detected all of those. The SNVs detected in other genes of the panel were not retained with our method for the following reasons: they were either identified as germline variants (n=5), detected in cancer but with insufficient coverage in non-cancer cells (n=3), detected but not in enough cancer cells (n=7), not detected despite sufficient coverage (n=3), or not covered (n=8) (**Supplementary Table S3**). Overall, 62% of the SNVs detected in the panel also found support in scRNA data. Since the scRNA-seq and panel-seq samples originated from different regions of the biopsy, the false negatives could be due to tumor heterogeneity or to the low VAF (<0.1) of some variants (n=4, **Supplementary Table S3**). In comparison, SComatic only detected one TP53 variant (P1) in LR scRNA-seq data. Of note, two deletions were found in panel sequencing, and they were detected manually in the LR scRNA-seq data (**Supplementary Table S3**).

### LongSom identifies subclones in LR scRNA-seq data matching subclones in scWGS data

LongSom detected 4 fusions in patient P1, 16 in P2, and 2 in P3, using CTAT-LR-fusion as described in (Qin et al. 2024) (**Supplementary Table S4**). Next, LongSom inferred the clonal structure of the tumors based on the SNVs and fusions it detected using BnpC. LongSom also inferred the clonal structure from CNA profiles in the same cells, using inferCNV (**Supplementary Figure S5, Methods**). We also clustered the cells based on their gene expression, manually annotated the cancer clusters, and used those clusters as transcriptomic validation. Finally, we used the subclones inferred from scWGS as external validation (**Figure 3)**.

In patient P1, LongSom found two cancer subclones based on SNVs and fusions, referred to as A and B (**Figure 6a**). The larger subclone A (n = 50 cells) was predominantly defined by a set of seven SNVs, including mtSNVs chrM:5179 and chrM:13635, and the smaller subclone B (n = 30 cells) was mainly defined by a set of seven SNVs, including mtSNV chrM:2815, as well as two fusions SMG7--CH507-513H4.1 and GS1-279B7.2--GNG4. In expression-based UMAP embedding, cancer cells formed two distinct expression clusters that near perfectly overlapped the genotypic cancer subclones found based on SNVs and fusions (**Figure 6 a-d**). Clonal assignments based on SNVs and fusions and on CNA data were also very similar (**Figure 6a,e, Supplementary Figure S5).** In patient P1’s matched scWGS data, we also found two aneuploid (cancer) subclones, CC1 and CC3, based on CNA profiles (**Supplementary Figure S2**). We found that 3 loci exclusively mutated in subclone A (chr7:25123800, chrM:5179,chrM:13635) were also exclusively mutated in subclone CC3, and one locus exclusively mutated in subclone B (chrM:2815) was exclusively mutated in subclone CC1 (**Figure 4e, 6a**). All subclone-specific loci had less than 5 reads coverage in scWGS clones, except for chrM loci, which had 186-861 reads coverage and were strictly subclone exclusive (**Figure 4e**). Therefore, using mtSNVs, we could confidently match scRNA-seq subclones to scWGS subclones.

**Figure 6:**
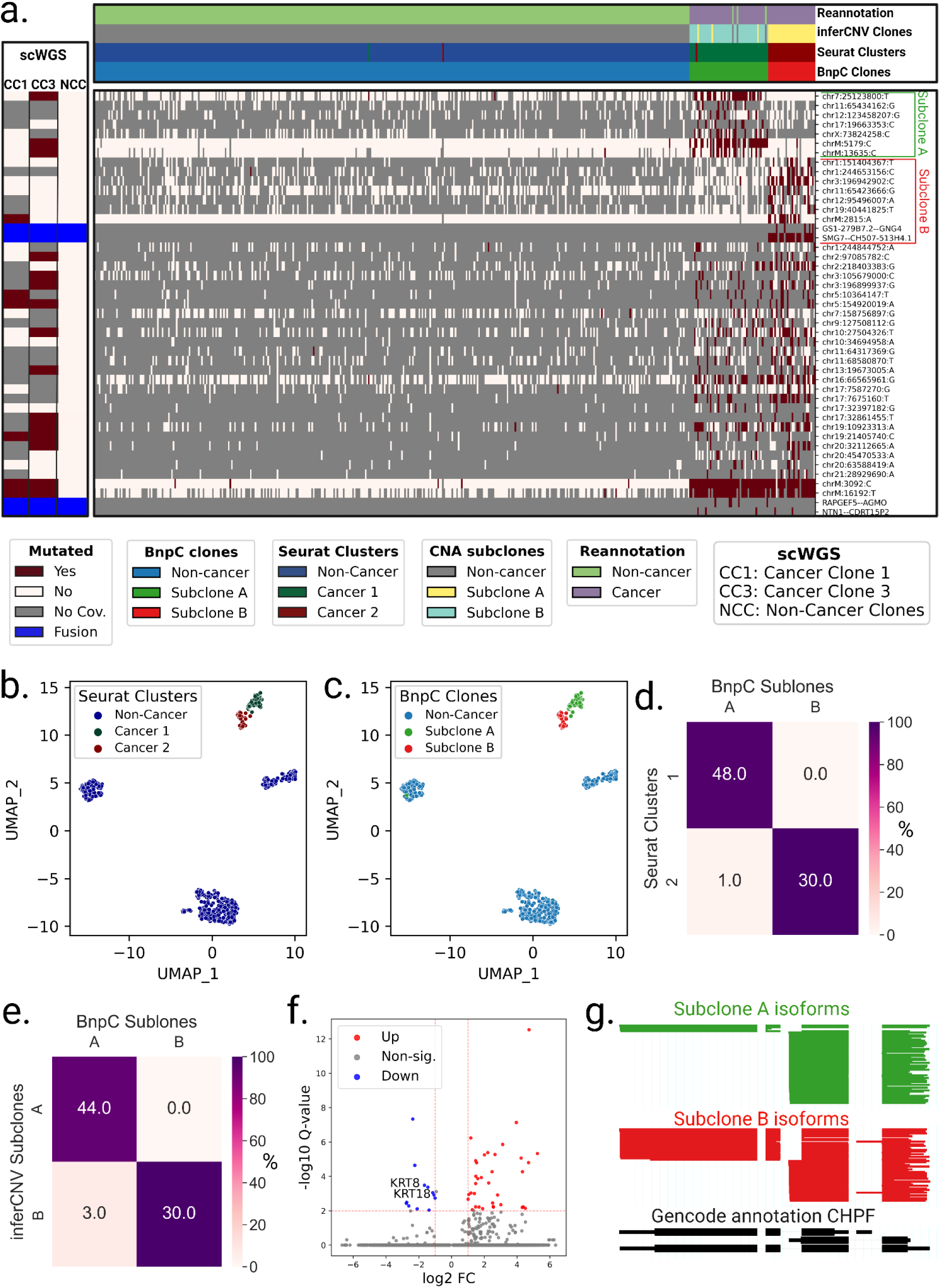
Analysis of intra-tumor heterogeneity using somatic variants detected in LR scRNA-seq in Patient P1. **a.** BnpC clustering of single cells from the tumor biopsy of patient P1 (columns) by somatic SNVs and fusions called by LongSom in LR scRNA-seq data (rows). Red indicates that a loci is mutated in a cell (beta-binomial P value < 0.05), white that it is not, and grey indicates no coverage in the cell at a given locus. Rows are colored according to the mutation status of aggregated scWGS diploid (Non-Cancer Clones) or aneuploid (Cancer Clone 1 and 3) cells. Fusions appear in blue. Columns are colored from top to bottom by cell types reannotated by LongSom, inferCNV CNAs subclones, expression clusters, and BnpC subclones inferred from somatic SNVs and fusions. **b,c.** UMAP embedding of patient P1 gene expression data, colored by (**b**) Seurat clusters and (**c**) BnpC subclones. **d,e.** Confusion matrix of cells in each expression-derived cancer cluster (rows) and **(d)** cells in the subclones inferred from BnpC, and **(e)** cells in the subclones inferred from inferCNV (columns), colored by the percentage of the total number of cells in each subclone and annotated by absolute numbers. **f**. Volcano plot of differentially expressed genes identified between subclones B and A. Keratin genes downregulated in subclone B are annotated. **g.** ScisorWiz representation of CHPF isoforms in subclones A and B. Colored areas are exons, whitespace areas are intronic space, not drawn to scale, and each horizontal line represents a single read colored according to subclones.

In patient P2, LongSom found one cancer clone using mutations and fusions, coinciding well with the aneuploid CNA scRNA clone and the gene-expression-based cancer cluster, similarly, in scWGS data we only saw one aneuploid CNA clone (**Supplementary Figure S2b, S5b, S6**). Therefore, all available data modalities point toward a monoclonal cancer population in this patient.

In patient P3, LongSom found one clone, coinciding with the gene expression-based cancer cluster, however, two aneuploid subclones were detected in both scWGS and scRNA-seq data using CNA analysis (**Supplementary Figures S2c, S5c, S7**). This difference could be due to the low number of cancer cells or inter-sample heterogeneity.

### Subclones identified in patient P1 have differing predicted treatment outcomes

To explore the potential therapeutic resistance of subclones A and B identified in patient P1, we investigated the genomic and transcriptomic variations between them. In subclone A, we identified a missense variant in the ferroptosis regulator *ALDH3A2* (Val321Leu, **Supplementary Table S5**) indicating a lower cisplatin resistance (Dong et al. 2023). In subclone B, we identified a missense in *CCAR2* (Arg722Trp, **Supplementary Table S5**), a suppressor of homologous recombination, indicating a potential resistance against PARP inhibitors (Iyer et al. 2022)). Therefore, based on SNVs, subclone A is more likely to be treatment-sensitive, while subclone B is more likely to be treatment-resistant. On the transcriptomic level, Subclone B had notably downregulated expression of keratin genes *KRT8* and *KRT18*, two epithelial markers used to differentiate HGSOC cells from non-cancer cells (**Figure 6f**, **Supplementary Figure S8a,b**). It has been shown in vitro that *KRT8* and *KRT18* have a protective effect against cell death (Bozza et al. 2018), and their losseads to increased invasiveness but also cisplatin sensitivity (Fortier et al. 2013). Subclone B is therefore more likely to be chemosensitive than subclone A. We additionally investigated differential isoform usage, and while both subclones were mostly similar, we found a significant difference in *CHPF* (**Figure 6g**), *MYL6*, the tumor suppressor *BTG2,* and *NUTM2B-AS1* (**Supplementary Figure S8c-e**), however, we could not predict their pathogenicity.

### LR greatly outperforms SR scRNA-seq for variants detection and clonal reconstruction

Finally, we aimed to compare LR to SR scRNA-seq ability to call somatic SNVs. The HGSOC study had LR and SR scRNA-seq data from the same cells available. When we applied SComatic to SR scRNA-seq, we found only 114 loci (7.3 times less than LongSom calls in LR data), with only 9 SNVs common to both technologies (**Figure 7a**). The lower amount of loci identified could be related to the fact that, while the SR dataset had 4.3 times more sequenced reads compared to LR (mean 117.4k vs. 26.9k reads per cell), it had 3.5 times fewer mapped bases (mean 11.4Gb mapped vs. 3.3Gb mapped) due to shorter read length (**Supplementary Figure S9a,b**). Notably, somatic SNVs identified in SR data contained 26% of 3’UTR variants, against 6% in LR data (Figure 3b, 7a). As reads are captured by their 3’ end poly(A), a large fraction of the SR coverage is located in 3’UTR as reads are too short to exceed it. This explains the over-representation of 3’UTR variants in SR data, and previous studies reported up to 40% of them (Muyas et al. 2024). To validate in scWGS data the somatic SNV loci identified in SR data, we pooled samples together due to the low number of calls in patients P1 and P3 (**Supplementary Figure S9c-e**). Out of the 18 loci called in SR scRNA-seq data with sufficient coverage in scWGS, 50% were supported as somatic in scWGS, a lower support than LR data, and no SNV supported as germline was found. Furthermore, SR data identified none of the panel-supported TP53 variants nor over clinically relevant variants. Therefore, we found fewer calls in SR data, and those had lower scWGS and bulk panel data support. Using those calls, we identified no clonal structure in either patient P1 or P3, and only identified a partial clonal reconstruction in P2, the patient in which we detected most SR calls (**Figure 7d, Supplementary Figure S9f-g**).

**Figure 7:**
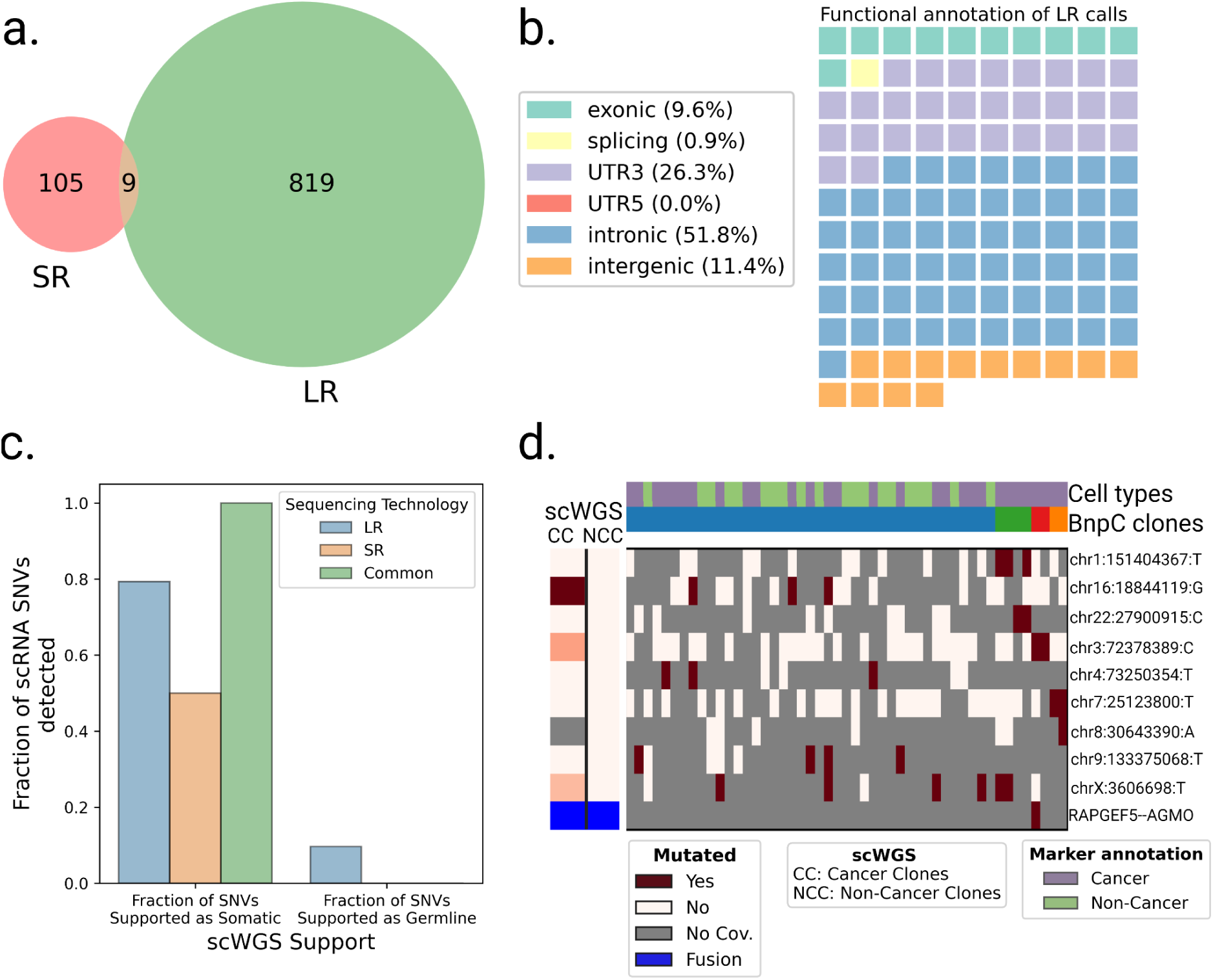
Analysis of SR scRNA-seq data and comparison with LR data. **a.** Venn diagram of the intersection between SR (SComatic) and LR (LongSom) somatic calls in LR scRNA-seq data from all patients aggregated. **b.** Waffle plot representing each of the 114 somatic SNVs detected by SComatic in SR scRNA-seq data, colored by their RefSeq functional annotation. **c.** Barplot of the fraction of all loci that is supported by scWGS data as either somatic or germline, colored by whether they were called in LR data, SR data, or both. **d.** BnpC clustering of single cells from the tumor biopsy of patient P1 (columns) by somatic SNVs and fusions (rows) called in SR scRNA-seq data. Red indicates that a loci is mutated in a cell (bet-binomial P value < 0.05), white that it is not, and grey indicates no coverage in the cell at a given locus. Rows are colored according to the mutation status of aggregated scWGS diploid (Non-Cancer Clones) or aneuploid (Cancer Clones) cells. Fusions appear in blue. Columns are colored from top to bottom by marker-based annotation and BnpC subclones inferred from somatic SNVs and fusions.

## Discussion

SNVs, mtSNVs, CNAs, fusions, gene expression, isoforms expression, and the micro-environment composition can all affect cancer treatment outcomes (Marine et al. 2020). Assessing all of these variations simultaneously from a single patient sample is particularly relevant in a clinical setting, where biological material is limited. Here, we show for the first time that this is possible using LR scRNA-seq data and we introduce LongSom, a workflow for detecting de novo somatic SNVs, fusions, and CNAs in LR scRNA-seq without matched normal. When applied to data from three HGSOC patients, we showed the Longsom outperformed SComatic for SNV detection and detected scWGS- and panel-validated SNVs, including clinically relevant ones. By integrating SNVs and fusions, LongSom successfully reconstructed the clonal heterogeneity and identified scWGS-matched subclones. Finally, in each subclone, we identified differentially expressed genes and subclone-specific SNVs with different implications for treatment resistance. Thus, we demonstrated that LR scRNA-seq is suitable for predicting treatment outcomes.

The performance of somatic SNV calling using non-cancer cells from the tumor biopsy as a “pseudo-normal” is contingent on reliable cell type annotations (Muyas et al. 2024). The cell-type reannotation step implemented in LongSom, based on the somatic variation profile of cells, led to the detection of up to 31% more somatic SNVs (patient P3) and significantly increased sensitivity without sacrificing precision.

LongSom is the first method combining de novo detection of SNVs, mtSNVs, and fusions from the same cell to reconstruct clonal heterogeneity. In the HGSOC dataset, the mitochondrial SNVs were called in most cancer and non-cancer cells, and some fusion calls were expressed in most clones or subclones (P2: IGF2BP2::TESPA1, P1: SMG7::CH507-513H4.1, etc.), making them ideal variations for cell-type reannotation and clustering. However, we demonstrated that mitochondrial SNVs require special filtering thresholds, as non-cancer cells frequently contained cancer mitochondrial reads. We showed that those reads were likely ambient mtRNA from dead or dying cancer cells encapsulated jointly with non-cancer cells during single-cell preparation. A conclusive answer to whether those are only technical artifacts or also originate from a biological mechanism, e.g. microenvironment revitalization (Liu et al. 2021; Zampieri et al. 2021), will require further investigation.

Despite the popularity of droplet-based scRNA-seq, multiple technical limitations remain unsolved, limiting the potential of downstream analysis. First, variant detection remains challenging due to the sparsity and low coverage of scRNA-seq assays, especially in LR assays despite rapid progress in the field (Dondi et al. 2023; Joglekar et al. 2023; Marx 2023; Al’Khafaji et al. 2024). To ensure that somatic calls are not germline polymorphisms, LongSom excludes sites with coverage <5 in non-cancer cells (39%), leading to potential false negatives. Second, read coverage is also uneven within a transcript, as transcripts produced by droplet-based scRNA-seq remain incomplete on the 5’ end due to intra-priming from intronic polyA-rich regions and on the 3’ end due to incomplete cDNA production (Nam et al. 2002; Hsu et al. 2022; Dondi et al. 2023; Verwilt et al. 2023). Third, RNA-seq is inherently limited to detecting only expressed SNVs and fusions. Nevertheless, as described above, LongSom detected a large fraction of variants in intronic or even intergenic regions. Last, similar to the mitochondrial reads contamination we observed, droplet-based scRNA-seq is sensitive to the encapsulation and subsequent sequencing of ambient RNA. This is especially true in cancer, where RNA from dead cancer cells is encapsulated with non-cancer cells, leading to false negative calls.

The performance of LongSom is dependent on a high sequencing quality (>Q20) and it has only been tested with PacBio data so far. Nanopore recently greatly improved its read quality and reached the Q20 threshold with chemistry R.10.4 (Ni et al. 2023), and LongSom will need to be tested on cancer biopsy scRNA-seq datasets generated with this technology.

In summary, LR scRNA-seq provides a unique snapshot of the cellular mechanisms by capturing multiple genomic and transcriptomic readouts from the same cell. With decreasing costs and increasing data size, we envision that LR scRNA-seq will become more common, potentially facilitating a better understanding of the processes underlying cancer treatment resistance. LongSom can be a valuable first step in guiding these analyses.

## Methods

### scRNA expression analysis

The raw sequencing data from HGSOC samples was retrieved from the European Genome-phenome Archive (EGA) with the accession number EGAS00001006807.

### Marker gene expression-based annotation

Cell annotation was retrieved from https://github.com/cbg-ethz/scIsoPrep/tree/master/bc_to_celltype (Dondi et al. 2023). We used “HGSOC” labels as cancer cells, and “Mesothelial.cells”, “Fibroblast”, “T.NK.cells”, “B.cells”, “Myeloid.cells”, “Endothelial.cells” labels as non-cancer cells.

### Clustering and visualization

Similar cells were grouped using Seurat FindClusters (Hao et al. 2024), and clusters with a majority (>90%) of non-cancer cells were grouped together as “non-cancer”. The results of the clustering and cell typing are visualized in a low-dimensional representation using Uniform Manifold Approximation and Projection (UMAP).

### Differential gene expression analysis

Differential expression was computed using Seurat FindMarkers (Hao et al. 2024), which uses a Wilcoxon test, corrected for multiple testing using the Bonferroni correction. A threshold of corrected P-value <0.01 and abs(log2(fold change))>1 was used for significance.

### Differential isoform usage analysis

Isoform classification and quantification were performed using scIsoPrep. Differential isoform testing was performed using a χ2 test as previously described in Scisorseqr (Joglekar et al. 2021). Differentially used isoforms were visualized using ScisorWiz (Stein et al. 2022).

### Somatic variants calling in LR scRNA-seq data with LongSom

To call somatic variants in LR scRNA-seq, we developed LongSom, a workflow implemented in python3 using Snakemake (Köster and Rahmann 2012) and available at https://github.com/cbg-ethz/LongSom. LongSom is designed to be run on a high-performance cluster. The HGSOC dataset was analyzed using 16 CPUs for a total of 64Gb memory for 3 hours.

### Preprocessing

PacBio long reads with minimal quality Q20 were de-concatenated, adapters were trimmed, demultiplexed, polyA tails were trimmed and finally, UMIs were deduplicated using scIsoPrep (https://github.com/cbg-ethz/scIsoPrep/tree/master) as described in (Dondi et al. 2023). Reads were mapped to the hg38 genome using minimap2 (Li 2018) with options -t 30 -ax splice -uf --secondary=no -C5.

### Error rates modeling

To distinguish true somatic SNVs from technical artifacts such as sequencing errors, mapping errors, or ambient RNA captured during cell encapsulation, LongSom models the background error rate using a beta-binomial distribution as described in (Muyas et al. 2024). Specifically, non-reference allele counts at homozygous reference sites are modeled using a binomial distribution with parameter P (error rate), which is a random variable that follows a beta distribution with parameters α and β, inferred using base count information from 500,000 sites in the genome randomly selected from patient P1 and P3 normal samples. Next, for each candidate loci, the beta-binomial distribution is used to test whether the non-reference allele counts are significantly higher than expected based on the error rate computed.

### Panel of normals

To discard positions affected by recurrent droplet-based scRNA-seq technical artifacts, LongSom uses a short-read PoN derived from (Muyas et al. 2024)). To also remove technical artifacts specific to LR-seq, we built an LR PoN using the two normal samples available from P1 and P3. The long-read PoN includes all sites with non-reference allele counts significantly higher than the background error rate modeled with the beta-binomial distribution in any of the normal samples provided. Matched samples were not included in the LR PoN during analysis: for P1, we only used the P3 normal sample and vice-versa.

### Cell-type re-annotation

To re-annotate cells, LongSom first identifies high-confidence cancer variants (HCCVs:, SNVs, mtSNVs and fusions). Candidate SNVs are identified using SComatic with default parameters except --min_mq 60. Then LongSom performes a series of eight filters: (1) loci with less than 20 reads coverage in aggregated cancer cells or aggregated non-cancer cells are filtered. (2) Longsom tests if, when ignoring the reads harboring candidate allele mutation reads, other non-reference allele counts at the locus are significantly higher than expected given the background error rate (beta-binomial test, significance threshold 0.05).. Loci (3) within homopolymers, (4) present in the gnomAD database (Chen et al. 2024) with a frequency of at least 1% of the total population, (5) present in RNA-editing databases (Tan et al. 2017, Kiran et al. 2012, Picardi et al. 2017), and (6) present in LR or SR PoN were filtered. (7) The SNVs where *VAF*_*NonCancer*_ < 0. 2, and (8) Δ*MCF* > 0. 4 are filtered, with Δ*MCF* defined as follows: Δ*MCF* = *MCF*_*Cancer*_ − *MCF*_*NonCancer*_ where *MCF* _(*Non*)*Cancer*_ is the fraction of mutated (non-)cancer cells, including only cells with minimum coverage of 1 at the position. Finally, (9) adjacent SNVs mapping within a 10,000 bp distance are filtered. SNVs passing all 8 filters are considered HCCVs. Forcing low *VAF* _*NonCancer*_ ensures that the candidate mutation is not a germline polymorphism or resulting from a loss of heterozygosity, while allowing *VAF*_*NonCancer*_ > 0 enables us to detect misannotated non-cancer cells. For samples with high levels of cross-contamination between cancer and non-cancer cells, higher *VAF*_*NonCancer*_ threshold can be used.

mtSNVs are considered HCCVs if they pass all SNV filters except (9) and (6). Instead, only the LR PoN is used. Fusions with a *MCF*_*Cancer*_ > 0. 05 and *MCF*_*NonCancer*_ < 0. 01 were selected as HCCVs.

Cells with less than 3 HCCV covered were filtered. Cells with at least 25% of covered HCCVs mutated were reannotated as cancer, while the others were reannotated as non-cancer.

### Somatic nuclear SNV identification

Candidate SNVs are identified using a modified version of SComatic (available at https://github.com/cbg-ethz/LongSom/tree/main/SComatic) with default parameters except --min_mq 60. LongSom then applies a set of 10 filters to identify somatic mutations, divided in three categories: coverage, noise, and germline filters. (1) loci with less than 5 reads coverage in aggregated cancer cells or aggregated non-cancer cells are filtered. (2) Loci with less than 3 alternative allele reads in at least 2 cancer cells are filtered. Then, LongSom applies filters intended to remove noise: (3) candidate somatic SNVs are distinguished from background noise and artifacts using a beta-binomial test parameterized using normal samples (Muyas et al. 2024), and loci with a non-significant test (threshold 0.001) in cancer cells were filtered as noise. (4) Longsom tests if, when ignoring the reads harboring candidate allele mutation reads, other non-reference allele counts at the locus are significantly higher than expected given the background error rate (beta-binomial test, significance threshold 0.05). (5) Mutations mapping within 4 bp or more of mononucleotide tracts (homopolymers) are filtered. (6) SNV loci present in RNA-editing databases (Tan et al. 2017, Kiran et al. 2012, Picardi et al. 2017) are filtered. (7) SNV loci present in either SR or LR panels of normals (PoN) are filtered. Finally, LongSom applies filters targeting germline variants: (8) Loci with a significant beta-binomial test in non-cancer cells (threshold 0.05) were filtered. Here, LongSom uses a stricter threshold than the original SComatic (0.001), in order to filter germline variants more efficiently. (9) SNV loci present in the gnomAD database (Chen et al. 2024) with a frequency of at least 1% of the total population were filtered. (10) Finally, adjacent SNV loci within a 10,000 bp distance are filtered, as these are likely misalignment artifacts in low-complexity regions or caused by allele-specific expression in cancer cells. This last filter was not part of the original SComatic.

### Somatic mtSNV calling

Due to the observed levels of cancer mitochondrial reads in non-cancer, we use specific rules to call somatic mtSNVs: (1) Loci with less than 100 reads coverage in aggregated cancer cells or aggregated non-cancer cells are filtered. (2) Loci present in the gnomAD database (Chen et al. 2024) with a frequency of at least 1% of the total population are filtered. (3) Loci where Δ*MCF* < 0.35 are filtered, with Δ*MCF* defined above. (4) Loci where *VAF*_*Cancer*_ < 0.1 are filtered.

Those *VAF* and Δ*MCF* parameters were determined based on the contamination level observed in the HGSOC dataset, and can be adjusted depending on the level of mitochondrial contamination.

### Somatic fusions identification

LongSom detects fusions on the single cell level using CTAT-LR-fusion v0.13.0 (https://github.com/TrinityCTAT/CTAT-LR-fusion/releases/tag/ctat-LR-fusion-v0.13.0) with standard options: -T fastq –vis (Qin et al. 2024). Fusions present in more than 5% of the cancer cells and less than 1% of the non-cancer cells were considered as somatic. LongSom allows fusions to appear in a low proportion of non-cancer cells as they can still harbor fusion reads due to ambient cancer RNA contamination.

### SNV annotation

SNVs were annotated using ANNOVAR (v2019Oct24) (Wang et al. 2010). An SNV was considered clinically relevant if it completed one of these conditions: it was flagged as pathogenic by ClinVar (Landrum et al. 2014), P-value was <0.05, the VEST (Carter et al. 2013) P-value was <0.05, the DANN (Quang et al. 2015) rankscore was <0.05, or FATHMM (Rogers et al. 2018) flagged it as deleterious.

### Single-cell genotyping

LongSom computes the alleles observed in each unique cell for each somatic SNV called. A cell is considered mutated at a position if the beta-binomial test is significant (with a significance threshold of 0.01) when applied to reads supporting the alternative allele compared to reads supporting the reference allele. For mtSNVs, to avoid false positives due to contamination, a cell is considered mutated if *VAF*_*Cell*_ > 0. 3, as determined from the HGSOC data (**Figure 4a,b**).

### Clonal detection based on SNVs and fusions

To detect subclones in cancer samples, LongSom only uses somatic SNVs covered in at least five cells, and fusions present in at least 3 cells, and then filters cells with less than three SNVs or fusions. Cells are then clustered using Bayesian non-parametric clustering (BnpC) (Borgsmüller et al. 2020), with arguments: cores (-n) 16, MCMC steps (--steps) 1000, alpha value of the Beta function used as prior for the concentration parameter of the Chinese Restaurant Process (--DPa_prior) [1,1], probability of updating the Chinese Restaurant Process concentration parameter (--conc_update_prob) 0, Beta(a, b) values of the Beta function used as parameter prior (--param_prior) [1,1].

### Clonal detection based on CNAs

LongSom first computes cell-gene matrices using featureCounts from Subread v2.0.6 (https://subread.sourceforge.net/) with parameters -L, using hg38 and gencode v36 as reference. It then uses those matrices as input for inferCNV to detect CNA subclones (https://github.com/broadinstitute/infercnv). For running CreateInfercnvObject, re-annotated non-cancer cells are used as a reference, and the parameter min_max_counts_per_cell=c(1e3,1e7) is used. For running inferCNV, the parameters cutoff=0.1 and leiden_resolution=0.01 are used. The CNA profiles displayed in this study are the ones obtained from the Hidden Markov Model learned by inferCNV.

### SNV calling using SComatic

As a comparison for LongSom, we called somatic SNVs in LR scRNA-seq using SComatic (Muyas et al. 2024). For this we used the marker-based cell types, and default parameters except the mapping quality (--min_mq) of 60 (maximum value for minimap2), the alpha and beta parameters computed for LR data (--alpha1 0.21, --beta1 104.95, --alpha2 0.25, --beta2 162.04), and a minimum distance (--min_distance) between loci of 0. We instead applied the same 10,000bp distance within “PASS” loci, as we did for LongSom.

### Empty droplets analysis

To estimate the mtSNV VAF in empty droplets, we first retrieved the empty droplets barcodes in each samples from the 10X Genomics CellRanger analysis. Then, we extracted the reads attached to those barcodes from the original bam file, and computed the VAF of chrM loci in each empty droplet barcode using pysam (version 0.21.0) pileup.

### scWGS

Cell suspensions were loaded and processed using the 10x Genomics Chromium platform with the single-cell CNV kit on the 10x Genomics Chromium Single Cell Controller (10x Genomics, cat. no. PN-120263) according to the manufacturer’s instructions. Paired-end sequencing was performed on the Illumina NovaSeq platform (100 cycles, 380 pm loading concentration with 1% addition of PhiX) at 0.1x depth per cell.

### Preprocessing and clonal reconstruction

Our scDNA-seq data analysis pipeline relied on CellRanger (https://www.10xgenomics.com/products/single-cell-cnv) for read mapping, quantification, binning, and GC and mappability correction. After filtering, we found 282 (P1), 182 (P2) and 290 (P3) cells per sample. The bin size used was 20kb, leading to very low coverage per bin. We further processed the resulting counts per bin to remove outlier bins with more than 3 times the median counts, and also outlier cells with highly imbalanced read count distribution as assessed by the Gini index. We used SCICoNE (Kuipers et al. 2020) to further segment the data into regions of at least 100 bins, resulting in CNAs that spanned at least 2Mb. We obtained subclonal copy number trees and assigned cells to the resulting CNA profiles. Subclones were considered as cancer subclones if they had an aneuploid CNA profile, and as non-cancer subclones if they had a fully diploid CNA profile.

### Variant allele calling in scWGS subclones

Cancer subclones were pooled together as well as non-cancer subclones due to low coverage (<10x per subclone). To determine if a loci was mutated in scWGS clones, we performed a beta binomial test parametrized on 500,000 sites randomly selected from aneuploid cells from all samples (significance threshold 0.01). We only considered loci called in scRNA-seq data for scWGS analysis if they fulfilled one of those two conditions: there was at least one mutated read in the locus or a least 17 reads coverage. Loci called in scRNA-seq data were considered as somatic if they were called in cancer clones only, and as germline if they were called in non-cancer clones.

### SNV validation in scWGS subclones

Cancer subclones were pooled together as well as non-cancer subclones due to low coverage (<10x coverage per subclone). To determine if a loci was mutated in scWGS clones, we performed a beta binomial test parametrized on 500,000 sites randomly selected from aneuploid cells from all samples (significance threshold 0.01). We only considered loci called in scRNA-seq data for scWGS analysis if they fulfilled one of two conditions: there was at least one mutated read in the locus or at least 17 reads coverage. Mutations called in scRNA-seq data were considered as somatic if they were called in cancer clones only, and as germline if they were called in non-cancer clones.

### De novo scWGS SNV calling

To call SNVs de novo in scWGS, we considered sites with at least 5 reads in both cancer and non-cancer cells in scRNA-seq, and with at least 5 reads in both aneuploid and diploid scWGS pooled clones (8-33M sites). Loci with at least 3 mutated reads in 2 aneuploid cells were considered. Loci with a beta-binomial test P < 0.001 in aneuploid clones and P>0.05 in diploid clones are considered somatic in scWGS. Sensitivity, precision and F1 performance statistics were computed as described in Muyas et al. 2024.

### Panel DNA sequencing

Panel sequencing was performed using the FoundationOne®CDx assay (FMI, Roche, Switzerland) (Milbury et al. 2022). DNA was extracted from FFPE tissue blocks with at least 20% tumor content with the Maxwell 16 FFPE Plus LEV DNA Purification Kit (AS1135, Promega, Dübendorf, Switzerland). Samples were assayed by adaptor ligation hybrid capture, performed for all coding exons of the FoundationOne panel. Sequencing was performed using the Illumina HiSeq instrument to a median exon coverage ≥500x.

## Data availability

All raw and processed sequencing data generated in this study have been submitted to the European Genome-phenome Archive (EGA, https://ega-archive.org/) under accession number EGASXXXXXXXXX.

## Software availability

LongSom is available at https://github.com/cbg-ethz/LongSom. All scripts necessary to reproduce this study are available at https://github.com/cbg-ethz/LongSom/tree/paper_version.

## Conflict of interest

The authors declare no competing interests.

## Acknowledgments

We thank Joanna Hård for her help with mitochondrial mutation contamination. We thank Ivan Topolsky for his support with cloud computing. We thank Lara Fuhrmann for naming LongSom. BioRender was used to generate the figures. A.D. and N.Bo were supported by the European Union’s Horizon 2020 research and innovation program under the Marie Sklodowska-Curie grant agreement (#766030 to N.Be). B.J.H. was supported by the National Cancer Institute grant U24CA180922.

## Author information

### Corresponding author

Correspondence to Niko Beerenwinkel

### Author contributions

N.Be. acquired the funding for this project. A.D. conceptualized the idea of this project. A.D. designed the LongSom pipeline with the help of N.Bo. The Tumor Profiler Consortium provided scWGS data and NGS panel results, and the full list of authors can be found in Supplementary Data S1. A.D. conducted all computational analyses except scWGS subclones inference (performed by P.F.). A.D. did all the visualizations and figures. A.D., N.Bo., F.J., and N.Be interpreted the results. A.D. and N.Bo. wrote the manuscript with contributions from all authors. B.J.H assisted with fusion transcript identification via CTAT-LR-fusion. All authors read and approved the final manuscript.

## Notes

### Competing Interest Statement

The authors have declared no competing interest.

### Summary of Updates

We implemented large changes to the SNV detection method, including more rigorous noise handling. All figures were updated. The manuscript was also restructured, and the mitochondrial contamination part was better introduced and demonstrated.

## Bibliography

Al’Khafaji AM, Smith JT, Garimella KV, Babadi M, Popic V, Sade-Feldman M, Gatzen M, Sarkizova S, Schwartz MA, Blaum EM, et al. 2024. High-throughput RNA isoform sequencing using programmed cDNA concatenation. Nat Biotechnol 42: 582–586.

Borgsmüller N, Bonet J, Marass F, Gonzalez-Perez A, Lopez-Bigas N, Beerenwinkel N. 2020. BnpC: Bayesian non-parametric clustering of single-cell mutation profiles. Bioinformatics 36: 4854–4859.

Bozza WP, Zhang Y, Zhang B. 2018. Cytokeratin 8/18 protects breast cancer cell lines from TRAIL-induced apoptosis. Oncotarget 9: 23264–23273.

Carter H, Douville C, Stenson PD, Cooper DN, Karchin R. 2013. Identifying Mendelian disease genes with the variant effect scoring tool. BMC Genomics 14 **Suppl 3**: S3.

Chen S, Francioli LC, Goodrich JK, Collins RL, Kanai M, Wang Q, Alföldi J, Watts NA, Vittal C, Gauthier LD, et al. 2024. A genomic mutational constraint map using variation in 76,156 human genomes. Nature 625: 92–100.

Dagogo-Jack I, Shaw AT. 2018. Tumour heterogeneity and resistance to cancer therapies. Nat Rev Clin Oncol 15: 81–94.

Dondi A, Lischetti U, Jacob F, Singer F, Borgsmüller N, Coelho R, Tumor Profiler Consortium, Heinzelmann-Schwarz V, Beisel C, Beerenwinkel N. 2023. Detection of isoforms and genomic alterations by high-throughput full-length single-cell RNA sequencing in ovarian cancer. Nat Commun 14: 7780.

Dong H, He L, Sun Q, Zhan J, Li J, Xiong X, Zhuang L, Wu S, Li Y, Yin C, et al. 2023. Inhibit ALDH3A2 reduce ovarian cancer cells survival via elevating ferroptosis sensitivity. Gene 876: 147515.

Foord C, Hsu J, Jarroux J, Hu W, Belchikov N, Pollard S, He Y, Joglekar A, Tilgner HU. 2023. The variables on RNA molecules: concert or cacophony? Answers in long-read sequencing. Nat Methods 20: 20–24.

Fortier A-M, Asselin E, Cadrin M. 2013. Keratin 8 and 18 loss in epithelial cancer cells increases collective cell migration and cisplatin sensitivity through claudin1 up-regulation. J Biol Chem 288: 11555–11571.

Gao R, Bai S, Henderson YC, Lin Y, Schalck A, Yan Y, Kumar T, Hu M, Sei E, Davis A, et al. 2021. Delineating copy number and clonal substructure in human tumors from single-cell transcriptomes. Nat Biotechnol 39: 599–608.

Gao T, Soldatov R, Sarkar H, Kurkiewicz A, Biederstedt E, Loh P-R, Kharchenko PV. 2023. Haplotype-aware analysis of somatic copy number variations from single-cell transcriptomes. Nat Biotechnol 41: 417–426.

Hao Y, Stuart T, Kowalski MH, Choudhary S, Hoffman P, Hartman A, Srivastava A, Molla G, Madad S, Fernandez-Granda C, et al. 2024. Dictionary learning for integrative, multimodal and scalable single-cell analysis. Nat Biotechnol 42: 293–304.

Hsu J, Jarroux J, Joglekar A, Romero JP, Nemec C, Reyes D, Royall A, He Y, Belchikov N, Leo K, et al. 2022. Comparing 10x Genomics single-cell 3’ and 5’ assay in short-and long-read sequencing. BioRxiv.

Iyer DR, Harada N, Clairmont C, Jiang L, Martignetti D, Nguyen H, He YJ, Chowdhury D, D’Andrea AD. 2022. CCAR2 functions downstream of the Shieldin complex to promote double-strand break end-joining. Proc Natl Acad Sci USA 119: e2214935119.

Jamal-Hanjani M, Quezada SA, Larkin J, Swanton C. 2015. Translational implications of tumor heterogeneity. Clin Cancer Res 21: 1258–1266.

Joglekar A, Foord C, Jarroux J, Pollard S, Tilgner HU. 2023. From words to complete phrases: insight into single-cell isoforms using short and long reads. Transcription 1–13.

Joglekar A, Prjibelski A, Mahfouz A, Collier P, Lin S, Schlusche AK, Marrocco J, Williams SR, Haase B, Hayes A, et al. 2021. A spatially resolved brain region- and cell type-specific isoform atlas of the postnatal mouse brain. Nat Commun 12: 463.

Kannan J, Mathews L, Wu Z, Young NS, Gao S. 2022. CAISC: A software to integrate copy number variations and single nucleotide mutations for genetic heterogeneity profiling and subclone detection by single-cell RNA sequencing. BMC Bioinformatics 23: 98.

Koshikawa N, Akimoto M, Hayashi J-I, Nagase H, Takenaga K. 2017. Association of predicted pathogenic mutations in mitochondrial ND genes with distant metastasis in NSCLC and colon cancer. Sci Rep 7: 15535.

Köster J, Rahmann S. 2012. Snakemake--a scalable bioinformatics workflow engine. Bioinformatics 28: 2520–2522.

Kramer CY. 1956. Extension of Multiple Range Tests to Group Means with Unequal Numbers of Replications. Biometrics 12: 307.

Kuipers J, Tuncel MA, Ferreira P, Jahn K, Beerenwinkel N. 2020. Single-cell copy number calling and event history reconstruction. BioRxiv.

Kwok AWC, Qiao C, Huang R, Sham M-H, Ho JWK, Huang Y. 2022. MQuad enables clonal substructure discovery using single cell mitochondrial variants. Nat Commun 13: 1205.

Landrum MJ, Lee JM, Riley GR, Jang W, Rubinstein WS, Church DM, Maglott DR. 2014. ClinVar: public archive of relationships among sequence variation and human phenotype. Nucleic Acids Res 42: D980–5.

Liu D, Gao Y, Liu J, Huang Y, Yin J, Feng Y, Shi L, Meloni BP, Zhang C, Zheng M, et al. 2021. Intercellular mitochondrial transfer as a means of tissue revitalization. Signal Transduct Target Ther 6: 65.

Li H. 2018. Minimap2: pairwise alignment for nucleotide sequences. Bioinformatics 34: 3094–3100.

Mansoori B, Mohammadi A, Davudian S, Shirjang S, Baradaran B. 2017. The different mechanisms of cancer drug resistance: A brief review. Adv Pharm Bull 7: 339–348.

Marine J-C, Dawson S-J, Dawson MA. 2020. Non-genetic mechanisms of therapeutic resistance in cancer. Nat Rev Cancer 20: 743–756.

Marx V. 2023. Method of the year: long-read sequencing. Nat Methods 20: 6–11.

Milbury CA, Creeden J, Yip W-K, Smith DL, Pattani V, Maxwell K, Sawchyn B, Gjoerup O, Meng W, Skoletsky J, et al. 2022. Clinical and analytical validation of FoundationOne®CDx, a comprehensive genomic profiling assay for solid tumors. PLoS ONE 17: e0264138.

Miller TE, Lareau CA, Verga JA, DePasquale EAK, Liu V, Ssozi D, Sandor K, Yin Y, Ludwig LS, El Farran CA, et al. 2022. Mitochondrial variant enrichment from high-throughput single-cell RNA sequencing resolves clonal populations. Nat Biotechnol 40: 1030–1034.

Morrison AC, Voorman A, Johnson AD, Liu X, Yu J, Li A, Muzny D, Yu F, Rice K, Zhu C, et al. 2013. Whole-genome sequence-based analysis of high-density lipoprotein cholesterol. Nat Genet 45: 899–901.

Mukherjee S, Bhatti GK, Chhabra R, Reddy PH, Bhatti JS. 2023. Targeting mitochondria as a potential therapeutic strategy against chemoresistance in cancer. Biomed Pharmacother 160: 114398.

Muyas F, Sauer CM, Valle-Inclán JE, Li R, Rahbari R, Mitchell TJ, Hormoz S, Cortés-Ciriano I. 2024. De novo detection of somatic mutations in high-throughput single-cell profiling data sets. Nat Biotechnol 42: 758–767.

Nam DK, Lee S, Zhou G, Cao X, Wang C, Clark T, Chen J, Rowley JD, Wang SM. 2002. Oligo(dT) primer generates a high frequency of truncated cDNAs through internal poly(A) priming during reverse transcription. Proc Natl Acad Sci USA 99: 6152–6156.

Ni Y, Liu X, Simeneh ZM, Yang M, Li R. 2023. Benchmarking of Nanopore R10.4 and R9.4.1 flow cells in single-cell whole-genome amplification and whole-genome shotgun sequencing. Comput Struct Biotechnol J 21: 2352–2364.

Osorio D, Cai JJ. 2021. Systematic determination of the mitochondrial proportion in human and mice tissues for single-cell RNA-sequencing data quality control. Bioinformatics 37: 963–967.

Qin Q, Popic V, Yu H, White E, Khorgade A, Shin A, Wienand K, Dondi A, Beerenwinkel N, Vazquez F, et al. 2024. CTAT-LR-fusion: accurate fusion transcript identification from long and short read isoform sequencing at bulk or single cell resolution. BioRxiv.

Quang D, Chen Y, Xie X. 2015. DANN: a deep learning approach for annotating the pathogenicity of genetic variants. Bioinformatics 31: 761–763.

Ramón Y Cajal S, Sesé M, Capdevila C, Aasen T, De Mattos-Arruda L, Diaz-Cano SJ, Hernández-Losa J, Castellví J. 2020. Clinical implications of intratumor heterogeneity: challenges and opportunities. J Mol Med 98: 161–177.

Rogers MF, Shihab HA, Mort M, Cooper DN, Gaunt TR, Campbell C. 2018. FATHMM-XF: accurate prediction of pathogenic point mutations via extended features. Bioinformatics 34: 511–513.

Serin Harmanci A, Harmanci AO, Zhou X. 2020. CaSpER identifies and visualizes CNV events by integrative analysis of single-cell or bulk RNA-sequencing data. Nat Commun 11: 89.

Shiau C-K, Lu L, Kieser R, Fukumura K, Pan T, Lin H-Y, Yang J, Tong EL, Lee G, Yan Y, et al. 2023. High throughput single cell long-read sequencing analyses of same-cell genotypes and phenotypes in human tumors. Nat Commun 14: 4124.

Stein AN, Joglekar A, Poon C-L, Tilgner HU. 2022. ScisorWiz: visualizing differential isoform expression in single-cell long-read data. Bioinformatics 38: 3474–3476.

Sun JX, He Y, Sanford E, Montesion M, Frampton GM, Vignot S, Soria J-C, Ross JS, Miller VA, Stephens PJ, et al. 2018. A computational approach to distinguish somatic vs. germline origin of genomic alterations from deep sequencing of cancer specimens without a matched normal. PLoS Comput Biol 14: e1005965.

Teer JK, Zhang Y, Chen L, Welsh EA, Cress WD, Eschrich SA, Berglund AE. 2017. Evaluating somatic tumor mutation detection without matched normal samples. Hum Genomics 11: 22.

Vasan N, Baselga J, Hyman DM. 2019. A view on drug resistance in cancer. Nature 575: 299–309.

Verwilt J, Mestdagh P, Vandesompele J. 2023. Artifacts and biases of the reverse transcription reaction in RNA sequencing. RNA 29: 889–897.

Wallace DC. 1994. Mitochondrial DNA sequence variation in human evolution and disease. Proc Natl Acad Sci USA 91: 8739–8746.

Wang K, Li M, Hakonarson H. 2010. ANNOVAR: functional annotation of genetic variants from high-throughput sequencing data. Nucleic Acids Res 38: e164.

Young MD, Behjati S. 2020. SoupX removes ambient RNA contamination from droplet-based single-cell RNA sequencing data. Gigascience 9: giaa151.

Zampieri LX, Silva-Almeida C, Rondeau JD, Sonveaux P. 2021. Mitochondrial transfer in cancer: A comprehensive review. Int J Mol Sci 22.

Zhang T, Jia H, Song T, Lv L, Gulhan DC, Wang H, Guo W, Xi R, Guo H, Shen N. 2023. De novo identification of expressed cancer somatic mutations from single-cell RNA sequencing data. Genome Med 15: 115.

Zhou Z, Xu B, Minn A, Zhang NR. 2020. DENDRO: genetic heterogeneity profiling and subclone detection by single-cell RNA sequencing. Genome Biol 21: 10.

