## Supplementary Figures S1-9 for "De novo detection of somatic variants in high-quality long-read single-cell RNA sequencing data"

### Supplementary Data

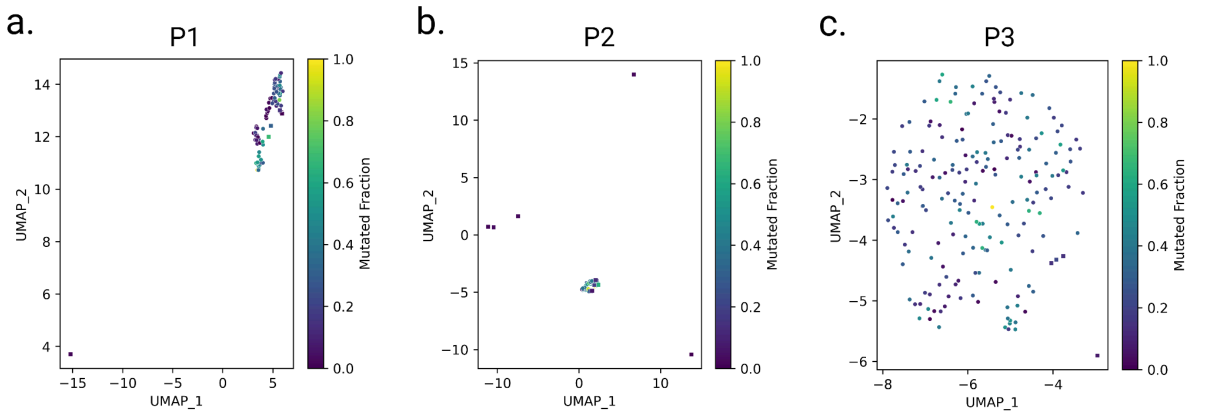

**Supplementary Figure S1: Mutational burden of re-annotated cells**

UMAP embeddings of LR scRNA-seq expression of cancer cells in patient **a.** P1, **b.** P2, **c.** P3. Cells are colored by the fraction of covered SNV loci that are mutated in the cell, and shaped by their re-annotation status, either previously annotated as cancer and re-annotated as cancer (circle) or previously annotated as non-cancer and annotated as cancer (square).

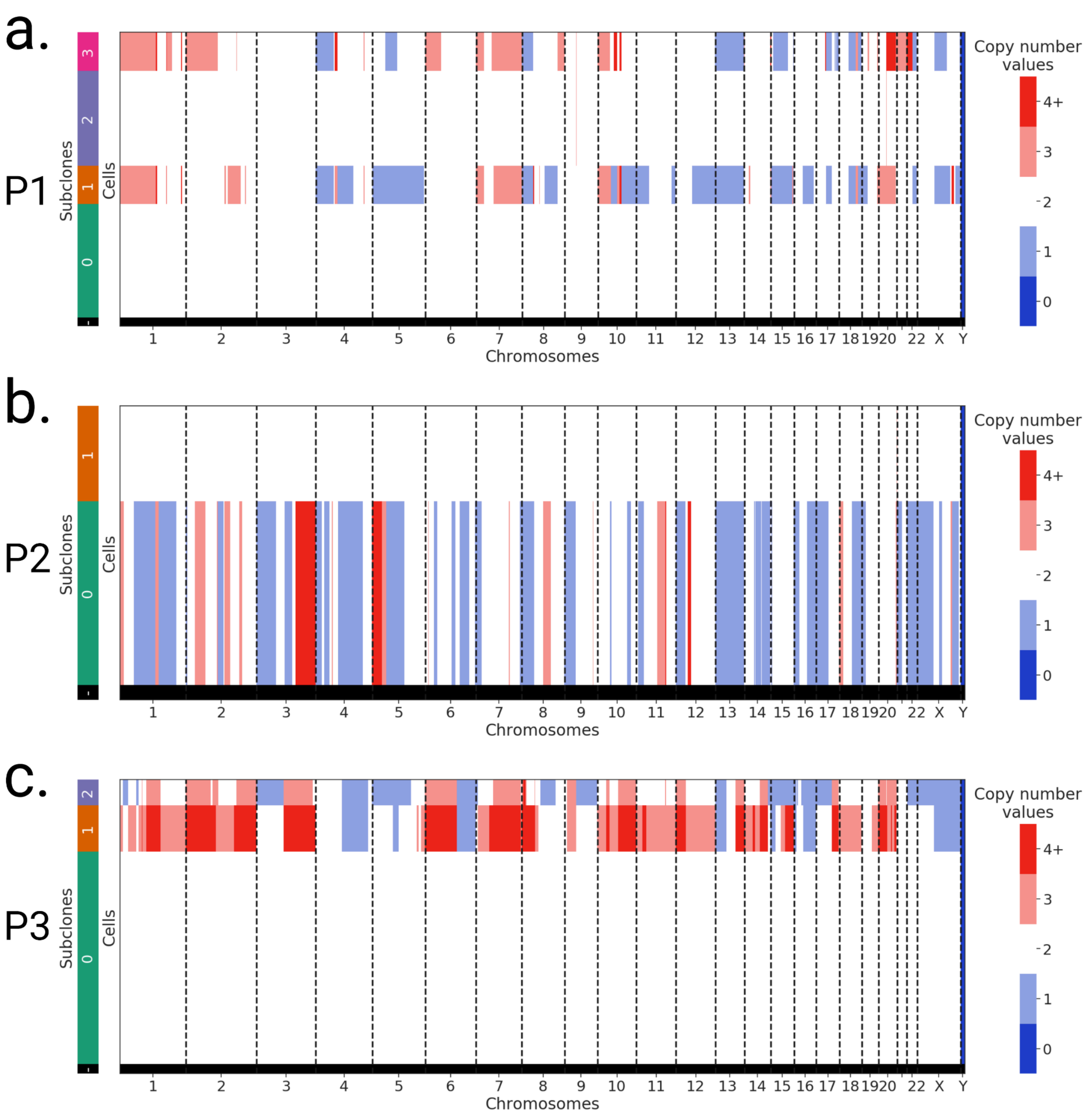

**Supplementary Figure S2: scWGS CNA subclones.**

scDNA-seq copy number values per subclone in **a.** patient P1, **b.** patient P2, and **c.** patient P3 data. Subclones with multiple copy number alterations are aneuploid (likely cancer), while copy number neutral subclones are diploid (likely non-cancer).

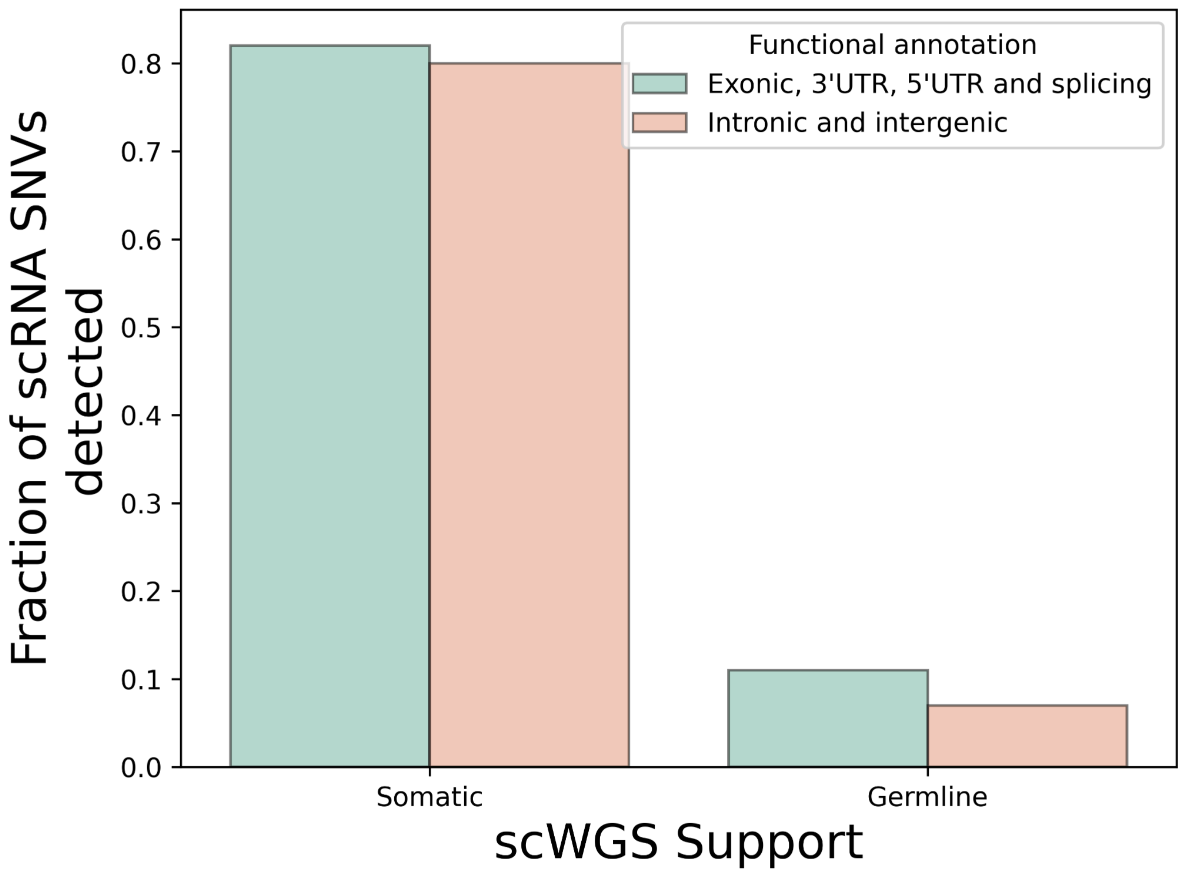

**Supplementary Figure S3: scWGS support depending on functional annotation**

Bar plot of the fraction of all loci being supported by scWGS data as either somatic or germline, colored by the functional annotation status of the loci.

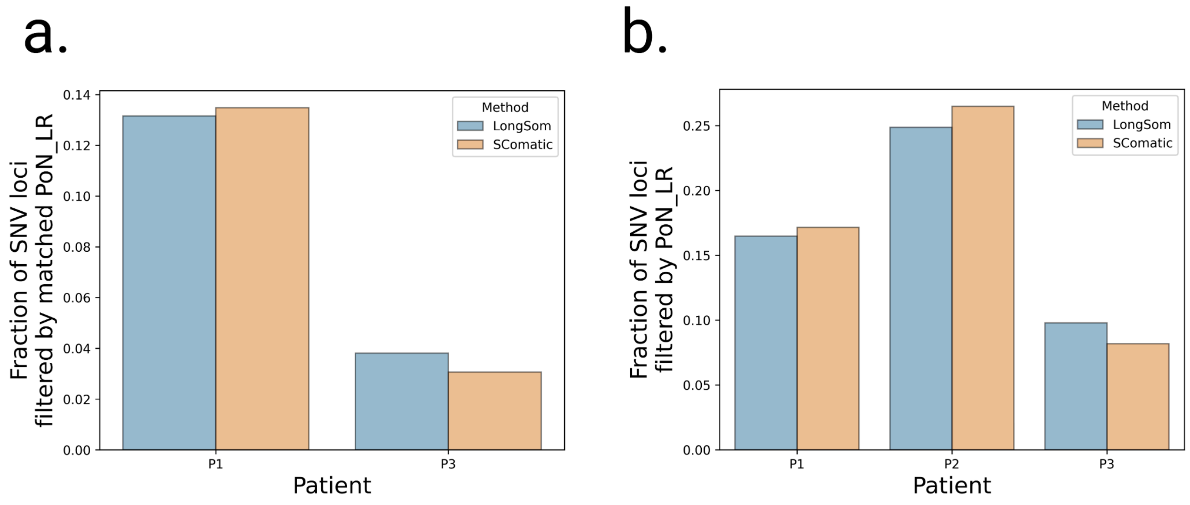

**Supplementary Figure S4: Validation using matching normals.**

**a.** Fraction of LongSom calls in somatic biopsies from patients P1 and P3 that were also called in matched normal samples **b.** Calls that were filtered with the LR PoN as the only filtering reason, as a fraction of the final LongSom call set.

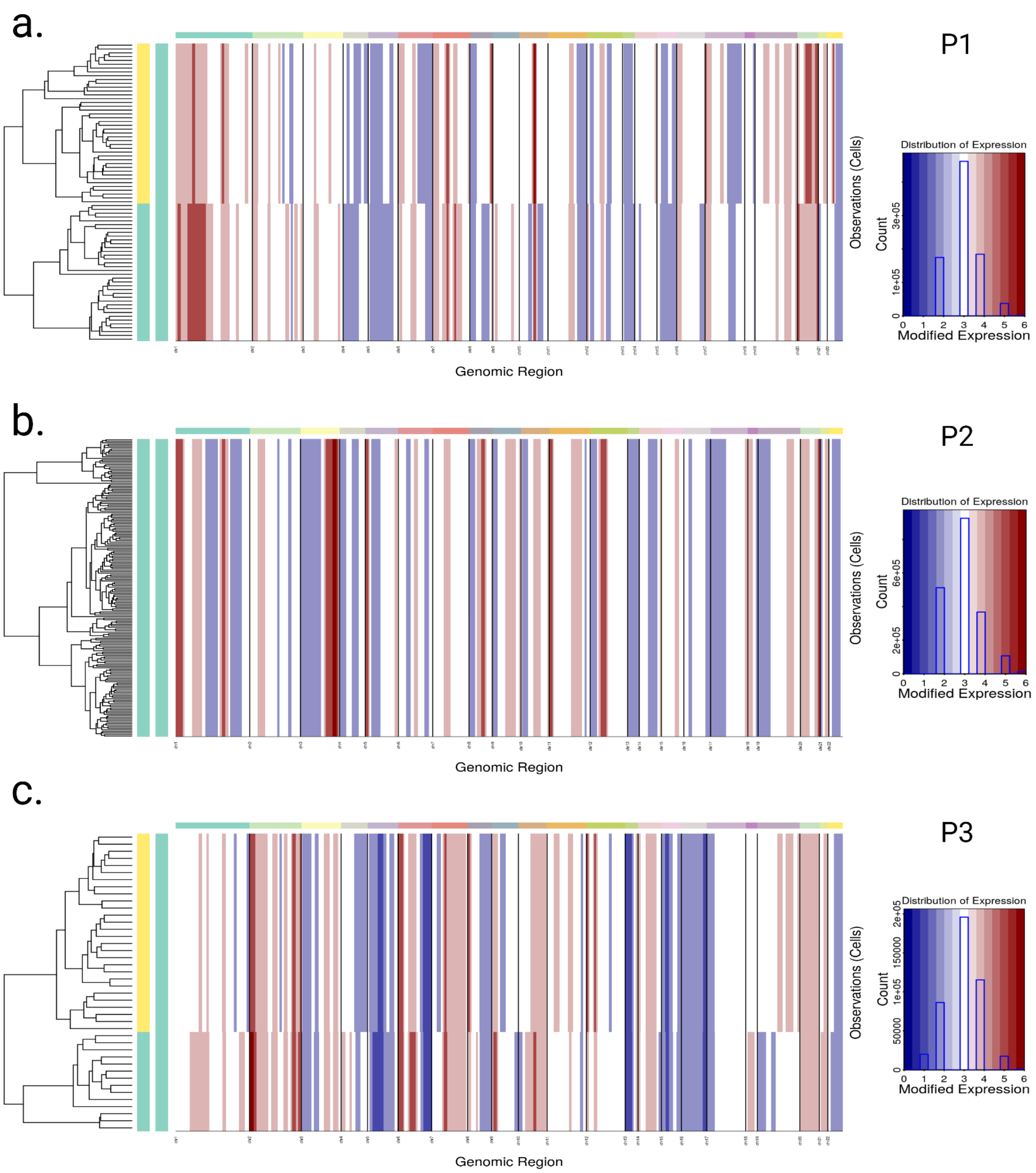

**Supplementary Figure S5: inferCNV CNA subclones.**

**a.** CNA profiles computed by inferCNV in patients **a.** P1, **b.** P2, and **c.** P3 cells by LongSom using inferCNV in LR scRNA-seq. Rows are individual cells, and colors (left) indicate subclonal attribution, columns are chromosomes. Regions in red have high copy number values, and in blue low.

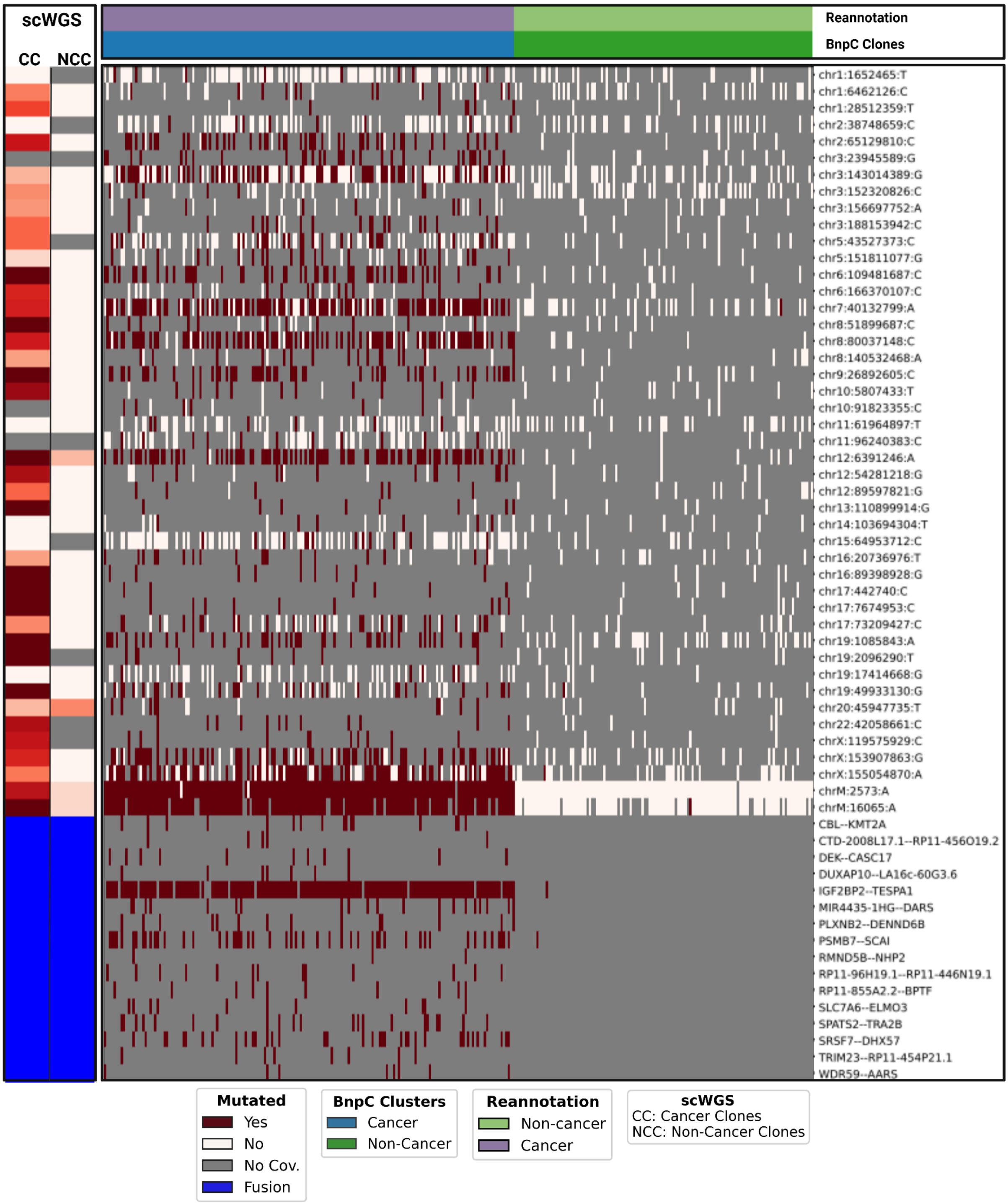

**Supplementary Figure S6: BnpC clonal reconstruction in Patient P2.**

BnpC clustering of single cells from the tumor biopsy of patient P2 (columns) by somatic SNVs and fusions called by LongSom in LR scRNA-seq data (rows). Red indicates that a loci is mutated in a cell (beta-binomial P value < 0.05), white that it is not, and grey indicates no coverage in the cell at a given locus. Rows are colored according to the mutation status of aggregated scWGS diploid (Non-Cancer Clones) or aneuploid (Cancer Clones) cells. Fusions appear in blue. Columns are colored from top to bottom by cell types reannotated by LongSom and BnpC subclones inferred from somatic SNVs and fusions.

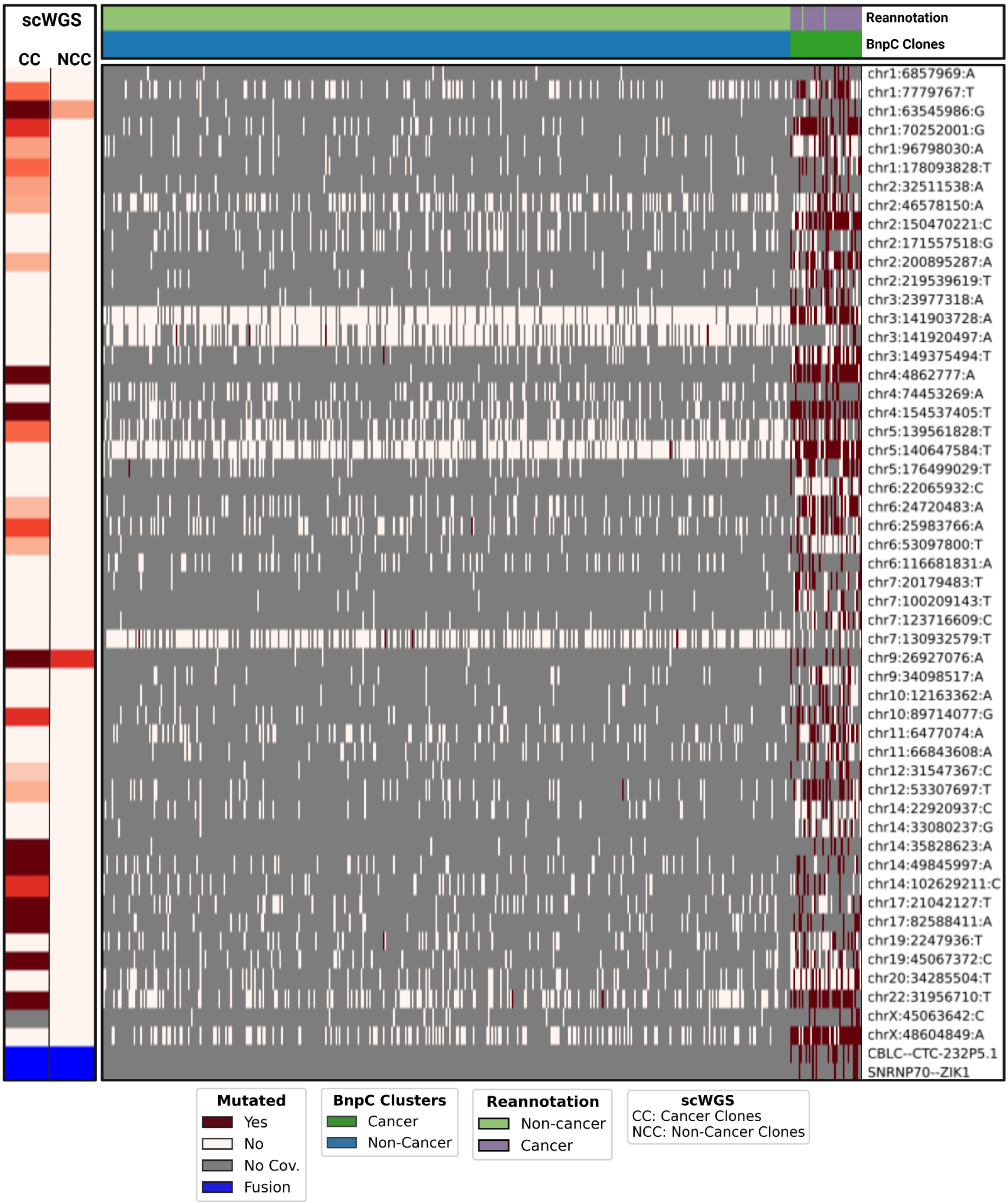

**Supplementary Figure S7: BnpC clonal reconstruction in Patient P3.**

BnpC clustering of single cells from the tumor biopsy of patient P3 (columns) by somatic SNVs and fusions called by LongSom in LR scRNA-seq data (rows). Red indicates that a loci is mutated in a cell (beta-binomial P value < 0.05), white that it is not, and grey indicates no coverage in the cell at a given locus. Rows are colored according to the mutation status of aggregated scWGS diploid (Non-Cancer Clones) or aneuploid (Cancer Clones) cells. Fusions appear in blue. Columns are colored from top to bottom by cell types reannotated by LongSom and BnpC subclones inferred from somatic SNVs and fusions.

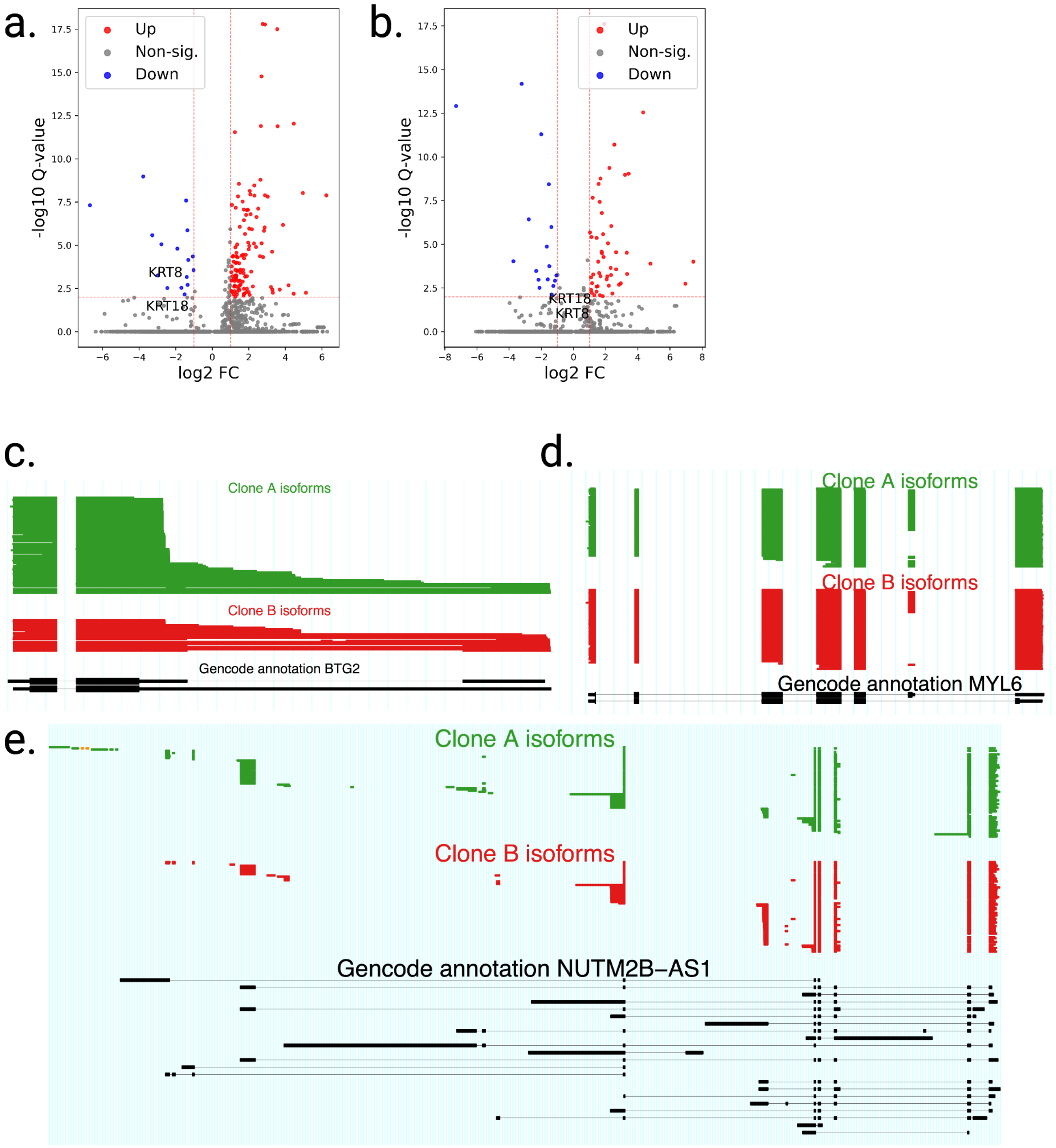

**Supplementary Figure S8: Differential gene and isoform expression in patient P1 subclones.**

**a**. Volcano plot of differentially expressed genes identified between Patient P1's subclone B and all other cancer subclones from all patients. **b.** Volcano plot of differentially expressed genes identified between Patient P1's subclone A and all other cancer subclones from all patients. **c.** ScisorWiz representation of *BTG2*, **d.** *MYL6* and **e.** *NUTM2B-AS1* isoforms. Colored areas are exons, whitespace areas are intronic space, not drawn to scale, and each horizontal line represents a single read colored according to subclones. Gencode notable reference isoforms appear in black.

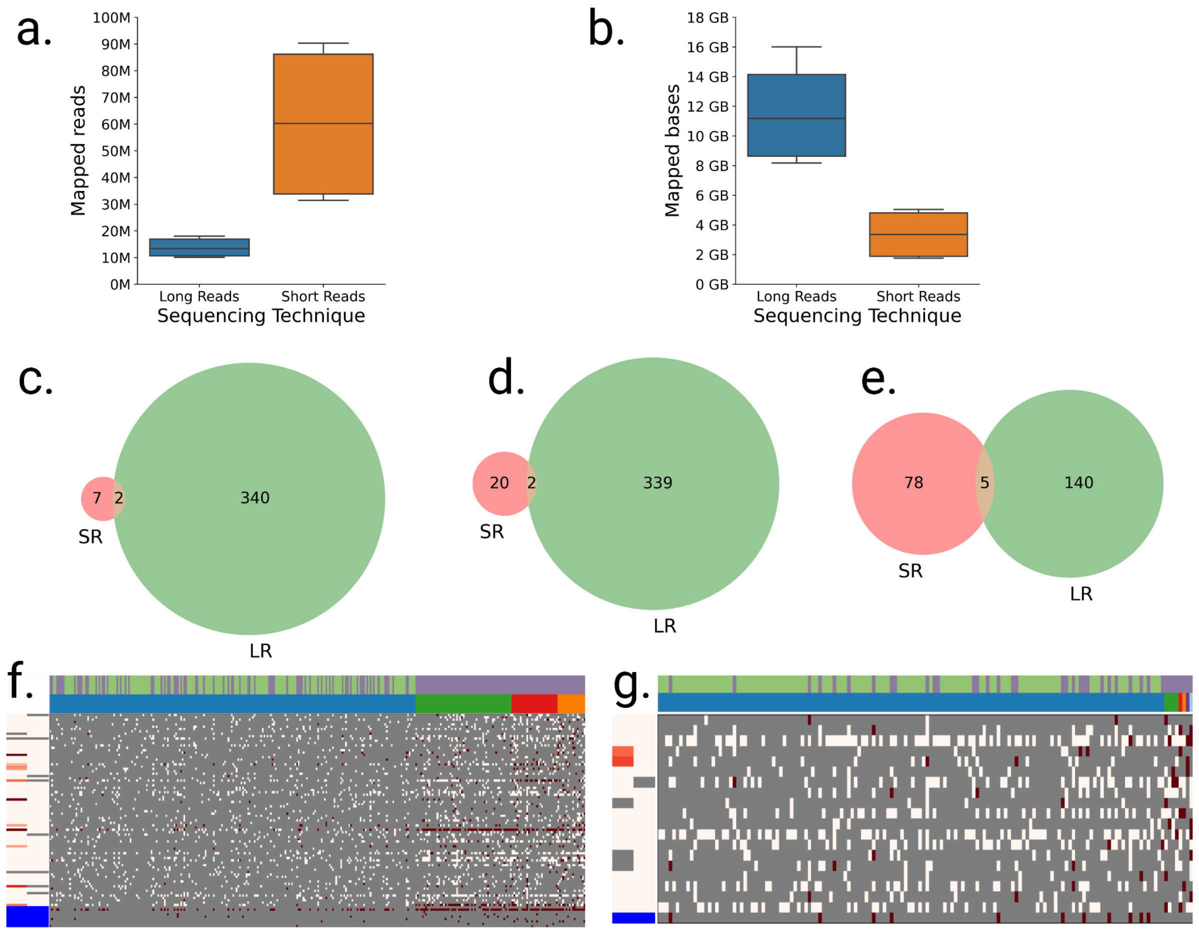

**Supplementary Figure S9: Sequencing statistics in LR and SR.**

**a.** Number of reads mapped across tumor and normal samples (n=5), per sequencing technique. **b.** Billions of bases mapped across tumor and normal samples (n=5), per sequencing technique. **c,d,e,** Venn diagram of the SNVs detected in scRNA-seq SR data by SComatic (red) and LR data by LongSom (green) in **c.** patient P1, **d.** patient P2 and **e.** patient P3. **f,g,** BnpC clustering of single cells (columns) from the tumor biopsy of patient **f.** P1 and **g.** P3 by somatic SNVs and fusions (rows) called in SR scRNA-seq data. Red indicates that a loci is mutated in a cell (bet-binomial P value < 0.05), white that it is not, and grey indicates no coverage in the cell at a given locus. Rows are colored according to the mutation status of aggregated scWGS diploid (Non-Cancer Clones) or aneuploid (Cancer Clones) cells. Fusions appear in blue. Columns are colored from top to bottom by marker-based annotation and BnpC subclones inferred from somatic SNVs and fusions.

**Supplementary Data S1: Tumor Profiler Consortium authors list**

Rudolf Aebersold[2], Melike Ak[28], Faisal S Al-Quaddoomi[9,17], Silvana I Albert[7], Jonas Albinus[7], Ilaria Alborelli[24], Sonali Andani[6,17,26,31], Per-Olof Attinger[11], Marina Bacac[16], Daniel Baumhoer[24], Beatrice Beck-Schimmer[39], Niko Beerenwinkel[4,17], Christian Beisel[4], Lara Bernasconi[27], Anne Bertolini[9,17], Bernd Bodenmiller[8,35], Ximena Bonilla[6,17,26], Lars Bosshard[9,17], Byron Calgua[24], Ruben Casanova[35], Stéphane Chevrier[35], Natalia Chicherova[9,17], Ricardo Coelho[18], Maya D'Costa[10], Esther Danenberg[37], Natalie Davidson[6,17,26], Monica-Andreea Drăgan[4], Reinhard Dummer[28], Stefanie Engler[35], Martin Erkens[14], Katja Eschbach[4], Cinzia Esposito[37], André Fedier[18], Pedro Ferreira[4], Joanna Ficek[6,17,26], Anja L Frei[31], Bruno Frey[13], Sandra Goetze[7], Linda Grob[9,17], Gabriele Gut[37], Detlef Günther[5], Martina Haberecker[31], Pirmin Haeuptle[1], Viola Heinzelmann-Schwarz[18,23], Sylvia Herter[16], Rene Holtackers[37], Tamara Huesser[16], Alexander Immer[6,12], Anja Irmisch[28], Francis Jacob[18], Andrea Jacobs[35], Tim M Jaeger[11], Katharina Jahn[4], Alva R James[6,17,26], Philip M Jermann[24], André Kahles[6,17,26], Abdullah Kahraman[17,31], Viktor H Koelzer[31], Werner Kuebler[25], Jack Kuipers[4,17], Christian P Kunze[22], Christian Kurzeder[21], Kjong-Van Lehmann[6,17,26], Mitchell Levesque[28], Ulrike Lischetti[18], Sebastian Lugert[10], Gerd Maass[13], Markus G Manz[30], Philipp Markolin[6,17,26], Martin Mehnert[7], Julien Mena[2], Julian M Metzler[29], Nicola Miglino[1], Emanuela S Milani[7], Holger Moch[31], Simone Muenst[24], Riccardo Murri[38], Charlotte KY Ng[24,34], Stefan Nicolet[24], Marta Nowak[31], Monica Nunez Lopez[18], Patrick GA Pedrioli[3], Lucas Pelkmans[37], Salvatore Piscuoglio[18,24], Michael Prummer[9,17], Natalie Rimmer[18], Mathilde Ritter[18], Christian Rommel[14], María L Rosano-González[9,17], Gunnar Rätsch[3,6,17,26], Natascha Santacroce[4], Jacobo Sarabia del Castillo[37], Ramona Schlenker[15], Petra C Schwalie[14], Severin Schwan[11], Tobias Schär[4], Gabriela Senti[27], Wenguang Shao[7], Franziska Singer[9,17], Sujana Sivapatham[35], Berend Snijder[2,17], Bettina Sobottka[31], Vipin T Sreedharan[9,17], Stefan Stark[6,17,26], Daniel J Stekhoven[9,17], Tanmay Tanna[4,6], Alexandre PA Theocharides[30], Tinu M Thomas[6,17,26], Markus Tolnay[24], Vinko Tosevski[16], Nora C Toussaint[9,17], Mustafa A Tuncel[4,17], Marina Tusup[28], Audrey Van Drogen[7], Marcus Vetter[20], Tatjana Vlajnic[24], Sandra Weber[27], Walter P Weber[19], Rebekka Wegmann[2], Michael Weller[33], Fabian Wendt[7], Norbert Wey[31], Andreas Wicki[30,36], Mattheus HE Wildschut[2,30], Bernd Wollscheid[7], Shuqing Yu[9,17], Johanna Ziegler[28], Marc Zimmermann[6,17,26], Martin Zoche[31], Gregor Zuend[32]

[1]Cantonal Hospital Baselland, Medical University Clinic, Rheinstrasse 26, 4410 Liestal, Switzerland, [2]ETH Zurich, Department of Biology, Institute of Molecular Systems Biology, Otto-Stern-Weg 3, 8093 Zurich, Switzerland, [3]ETH Zurich, Department of Biology, Wolfgang-Pauli-Strasse 27, 8093 Zurich, Switzerland, [4]ETH Zurich, Department of Biosystems Science and Engineering, Mattenstrasse 26, 4058 Basel, Switzerland, [5]ETH Zurich, Department of Chemistry and Applied Biosciences, Vladimir-Prelog-Weg 1-5/10, 8093 Zurich, Switzerland, [6]ETH Zurich, Department of Computer Science, Institute of Machine Learning, Universitätstrasse 6, 8092 Zurich, Switzerland, [7]ETH Zurich, Department of Health Sciences and Technology, Otto-Stern-Weg 3, 8093 Zurich, Switzerland, [8]ETH Zurich, Institute of Molecular Health Sciences, Otto-Stern-Weg 7, 8093 Zurich, Switzerland, [9]ETH Zurich, NEXUS Personalized Health Technologies, John-von-Neumann-Weg 9, 8093 Zurich, Switzerland, [10]F. Hoffmann-La Roche Ltd, Grenzacherstrasse 124, 4070 Basel, Switzerland, [11]F. Hoffmann-La Roche Ltd, Grenzacherstrasse 124, 4070 Basel, Switzerland, , [12]Max Planck ETH Center for Learning Systems, , [13]Roche Diagnostics GmbH, Nonnenwald 2, 82377 Penzberg, Germany, [14]Roche Pharmaceutical Research and Early

Development, Roche Innovation Center Basel, Grenzacherstrasse 124, 4070 Basel, Switzerland, [15]Roche Pharmaceutical Research and Early Development, Roche Innovation Center Munich, Roche Diagnostics GmbH, Nonnenwald 2, 82377 Penzberg, Germany, [16]Roche Pharmaceutical Research and Early Development, Roche Innovation Center Zurich, Wagistrasse 10, 8952 Schlieren, Switzerland, [17]SIB Swiss Institute of Bioinformatics, Lausanne, Switzerland, [18]University Hospital Basel and University of Basel, Department of Biomedicine, Hebelstrasse 20, 4031 Basel, Switzerland, [19]University Hospital Basel and University of Basel, Department of Surgery, Brustzentrum, Spitalstrasse 21, 4031 Basel, Switzerland, [20]University Hospital Basel, Brustzentrum & Tumorzentrum, Petersgraben 4, 4031 Basel, Switzerland, [21]University Hospital Basel, Brustzentrum, Spitalstrasse 21, 4031 Basel, Switzerland, [22]University Hospital Basel, Department of Information- and Communication Technology, Spitalstrasse 26, 4031 Basel, Switzerland, [23]University Hospital Basel, Gynecological Cancer Center, Spitalstrasse 21, 4031 Basel, Switzerland, [24]University Hospital Basel, Institute of Medical Genetics and Pathology, Schönbeinstrasse 40, 4031 Basel, Switzerland, [25]University Hospital Basel, Spitalstrasse 21/Petersgraben 4, 4031 Basel, Switzerland, [26]University Hospital Zurich, Biomedical Informatics, Schmelzbergstrasse 26, 8006 Zurich, Switzerland, [27]University Hospital Zurich, Clinical Trials Center, Rämistrasse 100, 8091 Zurich, Switzerland, [28]University Hospital Zurich, Department of Dermatology, Gloriastrasse 31, 8091 Zurich, Switzerland, [29]University Hospital Zurich, Department of Gynecology, Frauenklinikstrasse 10, 8091 Zurich, Switzerland, [30]University Hospital Zurich, Department of Medical Oncology and Hematology, Rämistrasse 100, 8091 Zurich, Switzerland, [31]University Hospital Zurich, Department of Pathology and Molecular Pathology, Schmelzbergstrasse 12, 8091 Zurich, Switzerland, [32]University Hospital Zurich, Rämistrasse 100, 8091 Zurich, Switzerland, [33]University Hospital and University of Zurich, Department of Neurology, Frauenklinikstrasse 26, 8091 Zurich, Switzerland, [34]University of Bern, Department of BioMedical Research, Murtenstrasse 35, 3008 Bern, Switzerland, [35]University of Zurich, Department of Quantitative Biomedicine, Winterthurerstrasse 190, 8057 Zurich, Switzerland, [36]University of Zurich, Faculty of Medicine, Zurich, Switzerland, [37]University of Zurich, Institute of Molecular Life Sciences, Winterthurerstrasse 190, 8057 Zurich, Switzerland, [38]University of Zurich, Services and Support for Science IT, Winterthurerstrasse 190, 8057 Zurich, Switzerland, [39]University of Zurich, VP Medicine, Künstlergasse 15, 8001 Zurich, Switzerland
